# A “*cooperative collaboration*” between HAND2 and MYCN shapes neuroblastoma cell identity

**DOI:** 10.1101/2022.05.18.492511

**Authors:** Man Xu, Ming Sun, Xiyuan Zhang, Rosa Nguyen, Haiyan Lei, Jack F. Shern, Carol J. Thiele, Zhihui Liu

## Abstract

Noradrenergic neuroblastoma (NB) is characterized by a core transcriptional regulatory circuitry (CRC) comprised of transcription factors (TFs) such as PHOX2B, HAND2 and GATA3, which form a network with MYCN. Physiological levels of MYCN mainly binds to promoters but aberrantly upregulated MYCN in NB also binds to enhancers. How MYCN invades enhancers, and what role the CRC TFs play in this process is unknown. Here we find that HAND2 assists MYCN to invade enhancers through an “*indirect cooperative*” TF-DNA binding mechanism. HAND2 cooperates with MYCN to compete with nucleosomes and regulate global gene transcription. The treatment of NB with an Aurora A kinase inhibitor plus HDAC inhibitor potently downregulates both MYCN and the CRC TFs and suppresses *MYCN*-amplified NB tumor growth. Our study identifies a “*cooperative collaboration*” among MYCN and CRC TFs and the simultaneous targeting these TFs is an effective way to treat this aggressive pediatric tumor.

## INTRODUCTION

Transcriptional regulation is typically mediated by distinct combinations of TFs, which work together to achieve the specific DNA binding that precisely regulates the gene transcription that controls developmental processes, lineage specification and function (1–5). Thus, it is not surprising that transcriptional dysregulation is a hallmark of many cancers as exemplified in the case of *MYC* family genes where their amplification and translocation disrupt normal developmental transcriptional programs (6–9). Recent studies have shown that under physiological conditions, MYC or MYCN transcription factor (TF) predominantly binds to promoter regions. However, in cancer cells, aberrantly elevated MYC or MYCN not only binds at gene promoters with canonical E-box motifs but also binds to enhancer regions containing low-affinity non-canonical E-boxes (10–12). This enhancer invasion is hypothesized to be one of the mechanisms by which MYC or MYCN drives tumorigenesis when it is aberrantly elevated. However, how MYC or MYCN invades enhancers is unknown.

Amplification of *MYCN* is found in poor prognostic neuroblastoma (NB), the most common extracranial solid tumor in children that derived from neural crest cells (13–15). During normal peripheral nervous system development, a set of TFs including PHOX2A, PHOX2B, HAND1, HAND2, GATA2, GATA3 and ASCL1 but not MYCN interacts as a network to determine sympathetic neuronal differentiation (14, 16–21). Interestingly, NB subverts a subset of these TFs including PHOX2B, HAND2, GATA3 and ASCL1 to form a core transcriptional regulatory circuitry (CRC) in noradrenergic NB (22–26). In NB cells, MYCN and CRC TFs PHOX2B, HAND2, GATA3, ISL1and TBX2 bind consensus DNA sequences adjacent to each other (24), suggesting that unlike in normal sympathetic neuronal cells, these sympathetic neuron fate determination TFs form a new network with the oncogene MYCN in NB. However, whether these CRC TFs and MYCN work together to achieve a specific DNA binding, or whether the CRC TFs facilitate MYCN to invade enhancers to initiate NB tumorigenesis or maintain the noradrenergic phenotype of NB cells remains unclear.

The concept of “*cooperative*” TF-DNA binding has evolved from decades of studies using *in vitro* cell free models, or cellular models focused on a few genes with some using reporter assays. Such studies identified that TF cooperativity arises through different mechanisms. “*Direct cooperative*” TF-DNA binding is dependent on protein–protein interactions between adjacently bound TFs, which may contribute to each other’s DNA binding, stabilize TFs on DNA to enhance their transcriptional activity (1,2,27–29). “*Indirect cooperative*” TF-DNA binding, also called “*collaborative*” DNA binding, occurs when motifs for each TF are separated by tens to hundreds of base pairs. These “*indirect cooperative*” TFs interactions do not require direct protein-protein interaction, but rather the TFs recognize adjacent binding sites within regulatory regions to compete with nucleosomes to access and cooperatively bind DNA (30–33). Importantly, through “*cooperative*” collaboration, the DNA binding of one TF helps the other TF to recognize new or low affinity binding sites. A limitation in the field is how well these models fit at a genome-wide level.

To interrogate the mechanisms of how CRC TFs and MYCN collaborates in NB at a genome-wide level, we used a combination of ChIP-seq, assay for Transposase-Accessible Chromatin using sequencing (ATAC-seq) and RNA sequencing (RNA-seq) assays. We demonstrate that the CRC TF HAND2, but not PHOX2B or GATA3 facilitates MYCN enhancer invasion. The cooperative binding of HAND2 and MYCN on DNA is accompanied by an increased chromatin accessibility, which leads to significant changes in global gene transcription. Our study uses genome-wide approaches to demonstrate the existence of “*cooperative*” TF-binding globally. More importantly our findings provide a rational for targeting these TFs simultaneously to increase the efficacy of cancer therapies. Indeed, the use of an Aurora A kinase inhibitor plus a HDAC inhibitor to target both MYCN and the CRC TFs, synergistically suppresses NB growth in a *MYCN*-dependent way. Here we discover the cooperativity between CRC TF HAND2 and MYCN at achieving specific DNA binding and global gene transcription regulation in NB and proposes targeting this “*cooperative collaboration*” through a rationale combined therapeutic approach.

## MATERIAL AND METHODS

### Cell culture

Human embryonic kidney cells (HEK293T) and Lenti-X HEK293 were obtained from ATCC and were maintained in Dulbecco’s modified Eagle’s media. Human neuroblastoma cell lines IMR32, IMR5, KCNR, LAN5, LAN6 and SK-N-FI were obtained from the cell line bank of the Pediatric Oncology Branch of the National Cancer Institute and have been genetically verified. All the NB cell lines were maintained in RPMI-1640 medium. All the cell culture medium was supplemented with 10%fetal calf serum (FBS), 100 μg/mL streptomycin, 100 U/mL penicillin, and 2 mM L-glutamine. Cells were grown at 37 °C with 5% CO2.

### Plasmids and stable clones

HA tagged MYCN (HA-MYCN) was generously provided by Dr. Wei Gu’s lab (34). HAND2 coding sequence tagged with 3x TY1 (TY1-HAND2) was synthesized by IDT company, then cloned into the doxycycline inducible pLVX-pTetOne-puro vector (Takara Bio) using In-Fusion HD (Takara Bio) following the manufacturer’s manual. To generate Hs_3_siHAND2 resistant HAND2 overexpression construct, HAND2 coding sequence with mutated nucleotides that cannot be recognized by Hs_3_siHAND2 while did not change the amino acid code was synthesized by IDT company. The mutant nucleotides in HAND2 are at the coding region that encodes amino acid 192 – 197 and the sequence can be found in Table S6. The HAND2 mutant (HAND2mut) open reading frame was cloned into the doxycycline inducible pLVX-pTetOne-puro vector (Takara Bio) using In-Fusion HD (Takara Bio) following the manufacturer’s manual. IMR32 cells were infected with lentiviral particles generated using either empty pLVX-TetOne-Puro vector or the pLVX-TetOne-Puro-HAND2mut vector, followed by puromycin selection. The stable clones named as IMR32tetEV and IMR32tetHAND2mut, respectively. HAND2 expression in IMR32tetHAND2mut could be induced with 0.1 μg/ml Dox treatment.

### CRISPR-Cas9 mediated gene knockout

To generate IMR32 cells that express the Cas9 protein, the cells were transduced with the lentivirus of Edit-R Inducible Lentiviral Cas9 (Dharmacon, Waltham, MA) followed with antibiotic selection by blasticidin. These cells express Cas9 when induced with Dox (0.5 μg/mL). Guide RNA (gRNA) that target the exon of human *HAND2* gene (sgHAND2) (Table S7) was designed using the online tool offered by Dharmacon. They were synthesized and cloned into the pLentiGuide-Puro vector (GenScript, Piscataway NJ) to be packaged into lentivirus using the Lenti-X HEK293 cells. The above mentioned Dox-inducible Cas9-expressing IMR32 cells were transduced with the lentivirus of sgHAND2 or non-targeting control gRNA (sgCtrl) and selected with puromycin. IMR32Cas9-sgCtrl or IMR32Cas9-sgHAND2 cell line was further single clone selected from the pool and cells are maintained in complete RPMI-1640 containing 0.5ug/ml puromycin. The depletion of *HAND2* is Dox (0.5 μg/ml) inducible.

### Transient transfection

Transient transfection was performed as described previously (35). siRNA controls and siRNAs targeting *HAND2, MYCN, TBX2, ISL1, PHOX2B* or *GATA3* were purchased from Qiagen or GE Dharmacon company (Table S7). siRNAs were transiently transfected into NB cells using Nucleofector electroporation (Lonza): solution L and program C-005 for IMR32 and IMR5; solution V and program A-030 for the rest NB cell lines. HA-MYCN or TY1-HAND2 plasmid was transfected into HEK293T cells using lipofectamine 2000.

To perform HAND2 knockdown rescue experiment, the Hs_3_siHAND2 that target *HAND2* was transiently transfected into either IMR32tetEV cells or IMR32tetHAND2mut cells. Right after transfection, the cells will be treated with or without Dox (0.1 μg/mL) to induce the expression of siHAND2 resistant form of HAND2.

### Cell growth and neurite extension assay

To evaluate cell proliferation, NB cells were plated in 96-well plates and the growth kinetics were monitored in IncuCyte ZOOM (Essen BioScience) using the integrated confluence algorithm as a surrogate for cell number. An alternative approach is to use CellTiter-Glo (Promega) to perform cell viability assay. Cell neurite length was measured using Essen IncuCyte ZOOM neurite analysis software. SynergyFinder online tool (https://synergyfinder.fimm.fi/synergy/20210322165414782814) was used to study the synergistic effect of the combination treatment of NB cells *in vitro*.

### Soft agar clonogenic assay

To assess effects of loss of *HAND2* on anchorage independent cell growth, 1 x 10^4^ IMR32Cas9-sgCtrl or IMR32Cas9-sgHAND2 cells were cultured in 0.7% top agarose in media on a layer of 1.4% bottom agar/media to prevent the adhesion of cells to the culture plates. Medium was changed twice a week with or without 0.5 μg/mL Dox, and visible colonies were observed after 2-4 weeks culture. The number of colonies was counted after crystal violet staining.

### Realtime PCR

The RNeasy Plus Mini Kit (Qiagen) was used to collect the mRNA according to the manufacturer’s protocol. Quantitative measurements of total β-actin and other genes’ levels were obtained using the BIO-RAD CFX Touch realtime PCR detection system and performed in triplicate. Ct values were standardized to β-actin level. Representative data from biological replicates were shown in this study. Primer sequences used for realtime PCR are shown in Table S6.

### Protein isolation, co-immunoprecipitation and western blotting analysis

For assessment of protein levels, cells were lysed using RIPA buffer, and 10 μg of total protein was separated and electroblotted as described previously (36). Protein bands probed with diluted primary antibodies (Table S8) were detected using a goat anti-rabbit or mouse IgG-HRP conjugated secondary antibody (200μg/mL; Santa Cruz Biotechnology) and visualized using enhanced chemiluminescence (Amersham Biosciences).

To detect the interaction between endogenous MYCN and the CRC TFs, co-immunoprecipitation (co-IP) was performed as previously described with slight modification (37). MYCN antibody (1 μg), or HAND2 antibody (1 μg), or normal IgG (1μg) was incubated with 50 μl Dynabeads M-280 sheep antimouse IgG magnetic beads (Thermo Fisher, 11202D) in 200ul wash buffer (50 mM pH 7.5 Tris-HCl, 137 mM NaCl, 1mM EDTA, 0.5% Triton X-100) overnight with rotation at 4 °C. IMR32 cells or NBLS cells were solubilized for 30 min in cold lysis buffer (50 mM pH 7.5 Tris-HCl, 137 mM NaCl, 1mM DTT, 1mM EDTA, 0.5% Triton X-100) supplemented with protease and phosphatase inhibitors (Halt protease and phosphatase inhibitor, Thermo), by shaking at 4°C. Whole cell extracts were clarified by a 10 min centrifugation at 20,000g at 4°C. The clear cell lysate (1 mg) was incubated with the Magnetic Beads coupled with MYCN antibody, HAND2 antibody or IgG control in total 1 ml lysis buffer and agitated at 4°C for 4 hr. Benzonase (500 U/ml) Mg2+ (2mM) will be added to the co-IP reaction if needed. Subsequently the beads were washed 5 times with washing buffer. The co-IP products were eluted by incubating with 25 μl 1x SDS loading buffer and boiling for 5 min. 5 μl of each sample was used for western blotting detection with10 μg cell lysate was used as Input. Primary antibodies of MYCN, HAND2, PHOX2B, GATA3, TBX2, ISL1 and MAX (Table S8) were used to detect the protein-protein interaction. To further investigate the interaction between MYCN with HAND2, HA-MYCN and TY1-HAND2 plasmids were co-transfected into HEK293T cells for 72h using Lipofectamine 2000. The cells were treated with Dox (0.5 μg/ml) to induce TY1-HAND2 expression. Similar co-IP experiments were performed as described above, while anti-TY1 antibody was used to pull down HAND2, anti-HA antibody was used to detect truncated MYCN protein.

### RNA-seq

Total RNA was isolated from IMR32 and LAN5 cells that have been transiently transfected with different siRNAs or siCtrl for 72 hr and subjected to RNA-seq analysis as previously described (38). Each sample had 3 biologic repeats. Total RNA was extracted using the RNeasy Plus Mini Kit (Qiagen Inc.) according to the manufacturer’s instructions. TruSeq® Stranded Total RNA LT Library Prep Kit (Illumina, San Diego, CA, USA) was used for preparing Strand-specific whole transcriptome sequencing libraries by following the manufacturer’s procedure. RNA-seq libraries were sequenced by paried-end with read length of 126 bp on Illumina HiSeq 2500, or HiSeq 3000/4000 of paired-end with read lenth of 150 bp. The Fastq files with 126bp or 150 bp paired-end reads were processed using Partek Flow. The raw reads are aligned using STAR and the aligned reads are quantified to annotation model through Partek E/M. The normalization method used here is counts per million (CPM) through Partek Flow. The statistic analysis of normalized counts used GSA or ANOVA. To get T-score, the normalized counts acquired from Partek Flow are exported and further analyzed using Parteck Genomics Suite v7.17. To eliminate batch effect, some of the CPM got from Partek Flow were analyzed using DESeq2. Statistical results of differentially expressed genes from Partek Flow, or Parteck Genomics Suite v7.17 or DESeq2 were analyzed using QIAGEN’s Ingenuity® Pathway Analysis (IPA®, QIAGEN) and gene set enrichment analysis (GSEA) (http://www.broadinstitute.org/gsea/index.jsp). By default, the false discovery rate (FDR) less than 0.25 is significant in GSEA.

### ATAC-seq

ATAC-seq was performed as previously described (38). IMR32 cells that have been transiently transfected with different siRNAs or siCtrl for 72 hr, or the IMR32tetHAND2mut cells have been treated with or without Dox for 36 hr to induce HAND2 were used for ATAC-seq. After digestion with trypsin, 100,000 viable cells were pelleted and the Tn5 transposition reaction was performed with the Nextera kit (Illumina), according to the manufacturer’s protocol. The amplified library was right side selected using SPRIselect reagent (Beckman catlog # B23317) following their instruction. ATAC libraries were sequenced on an Illumina NextSeq machine (2×75 cycles). The Fastq files with 75 bp paired-end reads were processed using Encode ATAC_DNase_pipelines (https://github.com/kundajelab/atac_dnase_pipelines) installed on the NIH biowulf cluster (https://hpc.nih.gov/apps/atac_dnase_pipelines.html).

### ChIP-seq

ChIP-seq was performed using the ChIP-IT High Sensitivity kit (Active Motif, cat. 53040) as described previously (38). Briefly, formaldehyde (1%, 15 minutes) fixed cells were sheared to achieve chromatin fragmented to a range of 200-700 bp using an Active Motif EpiShear Probe Sonicator. IMR32 cells that have been transiently transfected with different siRNAs or siCtrl for 24 hr or 72 hr, or the IMR32tetHAND2mut cells have been treated with or without Dox for 6 hr or 36 hr to induce HAND2 were used for ChIP-seq. IMR32 cells were sonicated at 25% amplitude, pulse for 20 seconds on and 30 seconds off for a total sonication “on” time of 16 minutes. Sheared chromatin samples were immunoprecipitated overnight at 4 °C with antibodies targeting MYCN, HAND2, PHOX2B, GATA3, H3K27ac, H3K4me4, H3K27me3, RNA Pol II and RNA Pol II S2P (Table S8). To normalize ChIP-seq signal, we employed Active Motif ChIP-seq spike-in using *Drosophila* chromatin (Active Motif catalogue # 53083) and an antibody against *Drosophila* specific histone variant H2Av (Active Motif, catalogue # 61686) according to the manufacturer’s instructions. ChIP-seq DNA libraries were prepared by Frederick National Laboratory for Cancer Research sequencing facility. Libraries were multiplexed and sequenced using TruSeq ChIP Samples Prep Kit (75 cycles), cat. # IP-2-2-1012/1024 on an Illumina NextSeq machine. 25,000,000-30,000,000 unique reads were generated per sample.

### ChIP-seq data processing

Previously published ChIP-seq datasets are down-loaded for this study, which including ChIP-seq datasets of MYCN, HAND2, PHOX2B, GATA3, TBX2 and ISL1 that generated in BE2C cells (GSE94822). For the home generated ChIP-seq data, ChIP enriched DNA reads were mapped to reference human genome (version hg19) using BWA. Duplicate reads were infrequent but discarded. For IGV sample track visualization, coverage density maps (tdf files) were generated by extending reads to the average size and counting the number of reads mapped to each 25 bp window using igvtools (https://www.broadinstitute.org/igv/igvtools).

ChIP-seq read density values were normalized per million mapped reads. High-confidence ChIP-seq peaks were called by MACS2 (https://github.com/taoliu/MACS) with the broad peak calling for H3K27me3 and RNA Pol II, narrow peak for TFs and the rest histone marks. The peaks which overlapped with the possible anomalous artifact regions (such as high-mappability regions or satellite repeats) blacklisted by the ENCODE consortium (https://sites.google.com/site/anshulkundaje/projects/blacklists) were removed using BEDTools. Peaks from ChIP-seq of MYCN, HAND2, PHOX2B, GATA3, H3K27ac, H3K4me3, H3K27me3, RNA Pol II, RNA Pol II S2P were selected at a stringent p-value (p<10^−5^ for PHOX2B and p<10^−7^ for the rest of the targets). Peaks within 1,000 bp to the nearest TSS were set as promoter. The distribution of peaks (as intronic, intergenic, exonic, etc.) was annotated using HOMER. Enrichment of known and *de novo* motifs were found using HOMER script “find Motifs Genome.pl” (http://homer.salk.edu/homer/ngs/peakMotifs.html).

The enhancers were identified using the ROSE2 (Rank Order of Super-Enhancers) software (https://github.com/BradnerLab/pipeline), using distal (>2,500 bp from TSS) H3K27ac peaks. Enhancer constituents were stitched together if clustered within a distance of 12.5 kb. The enhancers were classified into typical and super-enhancers based on a cutoff at the inflection point in the rank ordered set (where tangent slope = 1) of the ChIP-seq signal (input normalized) with p < 10^−7^.

ComputeMatrix function of the deepTools was used to generate a matrix of signal intensity of TFs of their peak centers (± 500 bp, total 1000 bp), as intensity scores in 10 bp bins. The matrix of signal intensity was further used to calculate the accumulated signal around each peak center to be used as signal intensity for each TF binding sites.

Heatmaps and composite plot of signal intensity of ChIP samples were generated using deepTools. Briefly, computeMatrix was used to calculate signal intensity scores per ChIP sample in a given genome region that was specified by a bed file. The output of computeMatrix was a matrix file of scores of two ChIP samples which was then used to generate the heatmap using plotHeatmap function, generate composite plot using plotProfile function.

### Monitoring of synergistic effects of drug combinations

The therapeutic effect of Aurora kinase A inhibitor (AURKAi) alisertib (MedChem Express, HY-10971) and HDACs inhibitor (HDACi) LBH589 (Medchem express, HY-10224) in MYCN-Amp NB cell lines IMR32, KCNR, IMR5 and MYCN single copy NB cell lines SH-N-FI, LAN6 was determined in a checkerboard fashion. Cell lines were seeded in two 96-well plates and incubated overnight. Each combination dose had 2 replications. Next day, cell lines were treated with different dose combination of alisertib and LBH589. Control cells were treated with DMSO. Each plate has their control cells. Cell viability was determined after 72 h using the CellTiter-Glo® luminescent assay (Promega, catalog number G9242). Cell viabilities of DMSO-treated cells were set to 100%. Results were graphed with GraphPad Prism software. IncuCyte® assay was used for testing the impact of synergistic effects of drug combinations to NB cell growth in realtime. Representative data from biological replicates were shown in this study.

### Xenograft Tumor Studies

To test the therapeutic effect of alisertib and LBH589 *in vivo*, 4-6 weeks old female athymic nude mice (Frederick Animal Facility, NCI) were orthotopically injected (injected through the adrenal fat pad into the adrenal gland) with 2.5×10^5^ IMR5-GFP-Luc cells (IMR5 cells stably transfected with a plasmid encodes both GFP and luciferase) into the in 30 μl Matrigel. Two weeks later, the luminescence signal in the tumor cells was measured by *in vivo* imaging system (IVIS) imaging. When the luminescence signal >1 x 10^8^, mice were grouped into 4 groups (10-11 mice/group): Group 1, treated with Vehicle; Group 2, treated with 10 mg/kg alisertib (disserved with 10% DMSO, 40% PEG300, 5% TWEEN-80, 45% saline) by oral gavage twice a day for 7 consecutive days (cycle days 1-7) in a 21-day cycle; Group 3, treated with 3 mg/kg LBH589 (disserved with 15% DMSO and saline) by intraperitoneal injection three times (M-W-F) a week; Group 4, treated with 10 mg/kg alisertib and 3 mg/kg LBH589 (see Fig. 7E schematic diagram for detailed information). After for 6 weeks (2 cycles) drug treatment, mice were euthanized, and tumors were collected and weighed. For the pharmacodynamics (PD) study, when the luminescence signal reached 1 x 10^8^ total flux [p/s], mice were grouped into 4 groups, which were treated with vehicle, alisertib, LBH589 and alisertib + LBH589, respectively. alisertib (10 mg/kg) was given at time 0 h, 8 h and 24 h, LBH589 (3 mg/kg) was given at time 0 h and 24 h. Mice were euthanized 4 hr after the final treatment and tumors were collected for western blot analysis. All Xenograft studies were approved by the National Cancer Institute’s Animal Care and Use Committee (ACUC), and all animal care was in accordance with institutional guidelines.

## RESULTS

### Association between CRC TFs and MYCN

To understand the cooperative nature of MYCN and the CRC TFs PHOX2B, HAND2, GATA3, ISL1, TBX2 on DNA binding and gene transcriptional regulation, we evaluated whether there is a physical interaction between MYCN and the other TFs. We performed a co-immunoprecipitation (co-IP) using an anti-MYCN antibody in *MYCN*-amplified IMR32 cell line followed by a western blot assay. The results showed that all the CRC components were isolated in the MYCN co-IP (Figure 1A). The interaction between MYCN and the CRC components, HAND2, PHOX2B, GATA3 and ISL1, was also observed in the NBLS (Figure S1A), a NB cell line that has a single copy of *MYCN* with relatively low MYCN protein levels. Of note, TBX2 was not detectable in NBLS cell line, which indicated that TBX2 was not a CRC component in this cell line. The treatment of samples with an endonuclease (benzonase) during co-IP completely blocked the interaction between MYCN and other TFs except for MAX (a known direct MYCN binding partner) and TBX2 (Figure 1A). This indicates that majority of these interactions are nucleic acid dependent. If there is cooperativity between MYCN and the CRC components on DNA binding during transcriptional regulation, this co-IP result suggests an “*indirect cooperative*” TF-DNA binding mechanism (28,30,31,39).

**Figure 1.**
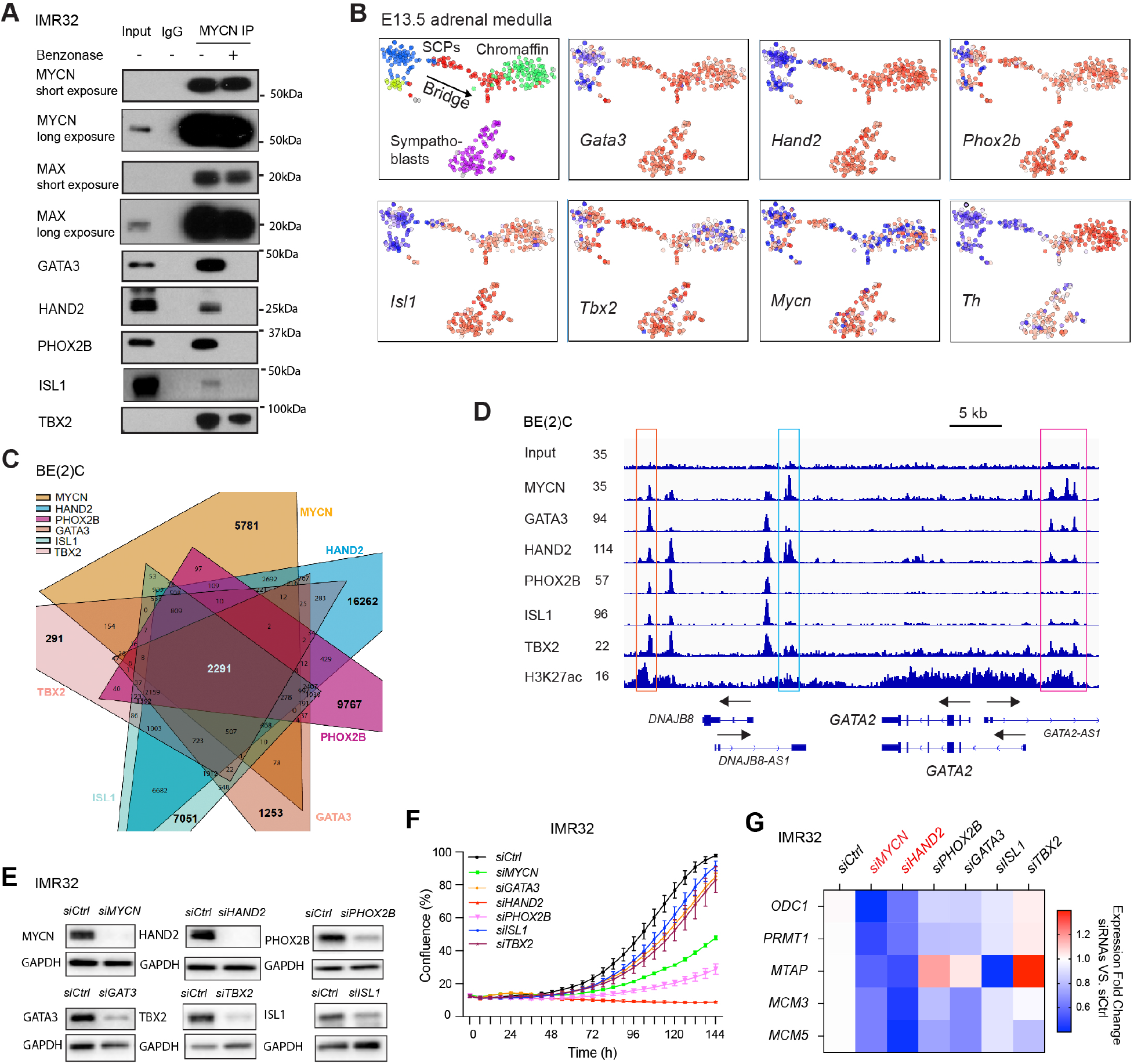
Association between NB CRC components and MYCN. (A) MYCN interacts with CRC TFs in IMR32 cells (*MYCN* amplified) shown by co-IP of MYCN protein followed by western blot. (B) The mRNA expression pattern of CRC components and *Mycn* in mouse embryonic adrenal medulla at single cell level is analyzed using Harvard interactive interface tools (http://pklab.med.harvard.edu/cgi-bin/R/rook/nc.SS2_16_250-2/index.html; single-cell RNA-seq results are available at Gene Expression Omnibus, GSE99933). Note of up-left corner image: SCPs, schwann precursor cells, blue color; Bridge, intermediate cell population undergo SCPs to chromaffin cell transition, red and yellow color; Chromaffin cells, green color; Sympathoblasts, purple color. Expression magnitude: blue lower, white average and red higher. Note of the rest images: low mRNA levels: blue; medium mRNA levels: white; high mRNA levels: red. (C) Intersection plot shows the overlap of ChIP-seq peaks of CRC components and MYCN determined by ChIP-seq data analysis (narrow peaks, p<1E-7). The combinatorial interaction and binding to the genomic DNA of these TFs could include all TFs or a few of them as a subgroup on certain genomic locus. (D) ChIP-seq signal tracks show the overlapped binding sites of CRC components and MYCN next to *GATA2* gene locus. Peaks within different color boxes indicate different patterns of co-localization among these TFs on different genomic loci. (E) The knockdown of each of the CRC components results in a decrease of their expression at protein levels shown by western blot assay. (F) The knockdown of CRC components results in a decrease of cell number shown by the IncuCyte cell confluence assay. (G) The regulation of a panel *MYCN* direct target genes by MYCN and the CRC TFs is detected by realtime PCR and shown in the heatmap.

To further study the correlation between *MYCN* and these CRC components, we investigated their expression pattern during normal development. Primary NB commonly arises along the sympathetic ganglia and in the medullary cells of the adrenal gland. The *MYCN* expression pattern in sympathoadrenal lineage is not well characterized but the noradrenergic NB CRC components *PHOX2B, HAND2* and *GATA3* are expressed in the sympathoadrenal lineage during development. A recent study in mouse embryos used single cell RNA sequencing (scRNA-seq) (20) to generate an atlas of gene expression in the developing adrenal medulla. By analyzing these scRNA-seq data, we found that *Phox2b, Gata3, Hand2, Isl1* and *Tbx2* are all highly expressed in sympathoblasts, in the intermediate stage cells (bridge) that arise from Schwann cell precursors (SCPs, which are late migrating neural crest cells), as well as in the chromaffin cells, but vary in SCPs (Figure 1B). Here *Th* serves as a chromaffin cell marker (Figure 1B). We found that *Mycn* mRNA levels are relatively high in sympathoblasts, but low in subsets of SCPs, bridge cells and chromaffin cells (Figure 1B). The scRNA-seq data showed that while CRC components are often co-expressed in most of the adrenal medulla cell populations, MYCN is co-expressed with the CRC TFs in the sympathoblasts but to a lesser degree in other cell types (Figure 1B).

We next compared the expression of CRC components and *MYCN* in human normal neural crest (NC), adrenal gland (AG) and neuroblastoma tumors (Versteeg cohort) by analyzing the R2 database. We found that the mRNA levels of these TFs were significantly higher in NB tumors than in either less differentiated NC cells or more differentiated mature adrenal gland derived cells (Figure S1B). This was consistent with the discovery that CRC components driven by super-enhancers were highly expressed in cancer cells (40). Of note, in some *MYCN*-nonamplified NB cell lines, *c-MYC* (or *MYCN*) was found to hijack the super-enhancer of *HAND2*, which drove the high expression of both *c-MYC* and *HAND2* (41). The scRNA-seq analysis of normal developing adrenal medulla showed that there appears to be variable to little overlap in expression of CRC components with *MYCN* in the bridge cells and early developing chromaffin cells (Figure 1B). However, all the CRC components and *MYCN* mRNA levels are high in NB (Figure S1B), suggesting NB cells may reflect an aberrant developmental state and the potential for different “combinatorial collaborations” among TFs in controlling gene regulation in NB compared to normal adrenal medulla. Given the findings that the *MYCN* transgenic mice and zebrafish form NB in the mouse adrenal gland or the equivalent tissue of zebrafish (42–44), these results suggest that the specific acquisition of high levels of *MYCN* in neuroblasts cooperates with the CRC TFs in driving NB tumorigenesis.

To assess the potential “*combinatorial collaborations*” among these TFs in NB, we analyzed the published ChIP-seq data of NB CRC components and MYCN in the SK-N-BE(2)C (BE(2)C) cell line (*MYCN* amplified) in detail (24). Using the Intervene tool (45), we identified both unique and overlapping binding sites among these TFs shown by the intersection plots of these TFs’ ChIP-seq peaks (Figure 1C). These overlaps suggest different combinatorial interactions among these 6 TFs. As an example, signal tracks showed different combinatorial co-localization among these TFs on genomic loci next to the *GATA2* gene locus (Figure 1D). The peaks within the orange box indicated a colocalization of all these 6 TFs on this locus while the peaks within the cyan box indicated the colocalization of MYCN and HAND2, and the peaks within the pink box show the colocalization of MYCN, GATA3, HAND2 and TBX2 (Figure 1D). Homer *de novo* motif scan of the unique peaks of each TF identified canonical binding motifs for these TFs (Figure S1C, left panel). The HAND2 binding motif matches the one for TWIST1, which is a non-canonical enhancer box (E-box) that is known to be a binding motif for many of the bHLH TFs. Moreover, HAND2 has two characterized binding motifs, one is non-canonical E-box (CATCTG) (22) and the other one is CTGGTTTTTGTC (derived from GSE61475). Interestingly, when focused on MYCN ChIP-seq peaks that overlapped with all the binding sites of the CRC components (2291 peaks), we found that the top ranked motifs included PHOX2A/2B, GATA3, ISL1, HAND2 and TCF4 binding motifs (Figure S1C, right panel). The TCF4 motif is a non-canonical E-box that is also a known binding motif for bHLH TFs. Similar results were observed by focusing on the TF ChIP-seq peaks that overlapped with the other TFs, where the top ranked peaks include their own binding motifs, as well as the motifs of the other TFs (Figure S1D). The motif scan result was consistent with the CRC component ChIP-seq results showing these TFs bind to consensus sequences and close to each other (24). The canonical E-box (CACGTG), a high-affinity MYC binding motif (46) were enriched in MYCN unique peaks while a MYCN low-affinity binding motif of the non-canonical E-box (CANNTG) was observed in the ChIP-seq peaks of each of the TFs that had overlapping binding sites (Figure S1C,D). Unlike MYCN, the binding motif of most of the CRC components identified in the unique peaks was also found in the overlapped peaks. These results suggest that the binding of MYCN at the overlapped binding sites with low-affinity E-boxes or other motifs might need to be assisted by the other TFs or might require the presence of the other TFs.

To investigate the biological function of the CRC components and the neural crest development related TFs that interact with MYCN in NB, we first mined the data from Project Achilles genome-wide CRISPR-Cas9 screen of a panel of pediatric cancer cell lines, and the results showed that most NB cell lines are preferentially dependent on these TFs (Figure S1E), which is consistent with essential role of these TFs identified by others (22–25). We also silenced each of these TFs using small interference RNAs (siRNAs) in IMR32 cells and found that the loss of these TFs (Figure 1E) resulted in a significant decrease in cell number based on IncuCyte cell confluence assay (Figure 1F). A model consistent with *cooperative binding* between a CRC component and MYCN would predict that a TF should work together with MYCN to regulate the expression of a common subset of MYCN target genes. We picked five well known MYCN direct target genes and performed realtime PCR to investigate how they were regulated after knocking down each of these TFs. In this panel of MYCN targets, the transcriptional activity of HAND2 most closely phenocopied MYCN (Figure 1G). Of note, GATA3, PHOX2B and HAND2, but not TBX2 or ISL1 have been broadly identified as noradrenergic CRC TFs in the tested NB cell lines (22–24). Therefore, we selected HAND2 as a candidate to interrogate the potential cooperativity between CRC components and MYCN at the genome-wide level of DNA binding and regulation of gene transcription.

### CRC component *HAND2* regulates the expression of *MYCN* targets and is essential in NB

To confirm the interaction between HAND2 and MYCN, we co-transfected TY1-tagged HAND2 and HA-tagged MYCN constructs into 293T cells followed by co-IP using an anti-TY1 antibody and found that MYCN was pulled down by HAND2 (Figure S2A). Co-IP of endogenous HAND2 in IMR32 cell lysates using an anti-HAND2 antibody also pulled down MYCN, supporting an interaction between HAND2 and MYCN (Figure S2B).

We extended the biological function analysis of *HAND2* by knocking down its expression in additional NB cell lines (Figure S2C). We observed that the silencing of *HAND2* significantly reduced viable cell number in all these tested NB cell lines (Figure 2A). To confirm that the biologic function was not due to an off-target effect of *HAND2* siRNA, we performed a rescue experiment. We generated a doxycycline (Dox) inducible *HAND2* overexpression IMR32 stable clone, in which the plasmid that encodes *HAND2* was mutated so that the siRNA did not recognize the nucleic acid coding sequence, but the amino acid codon was unchanged. We found that the Dox treatment induced overexpression of the *HAND2* mutant attenuated the *siHAND2* induced upregulation of *GAP43* mRNA levels and repression of cell proliferation (see Method section, Figure S2D, E).

**Figure 2.**
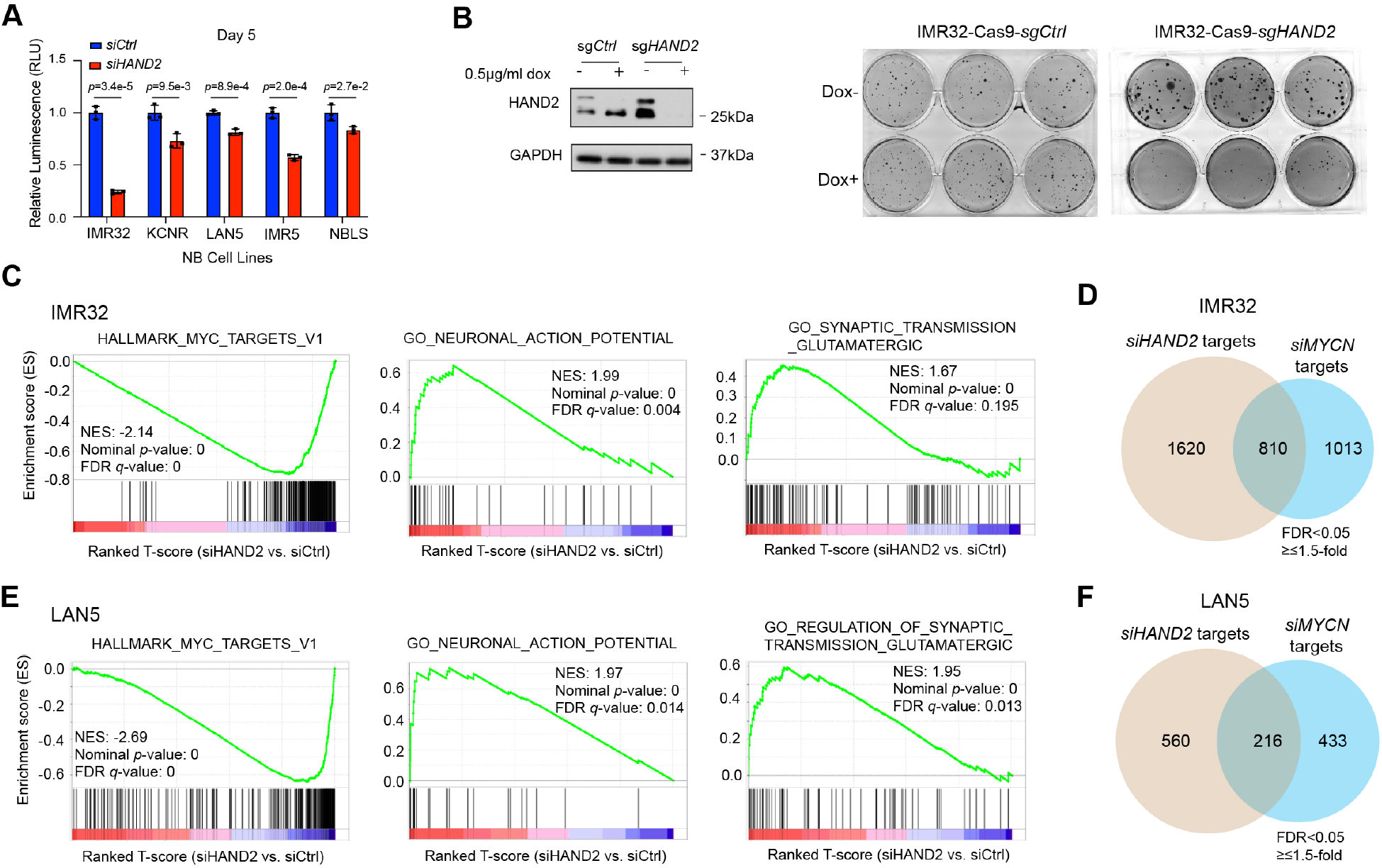
CRC component *HAND2* regulates the expression of *MYCN* targets and is essential in NB. (A) The knockdown of *HAND2* using siRNA results in a decrease of cell proliferation in multiple NB cell lines shown by CellTiter-Glo assay. The *p*-value indicated is calculated in Student’s t-Test. (B) Depletion of *HAND2* through doxycycline (Dox) inducible Crispr/Cas9 expression and a single-guide RNA (sgRNA) that targets *HAND2* but not control (sgCtrl) results in a decrease of HAND2 at protein levels after 7 days Dox treatment (left panel), which results in a decrease of anchorage-independent cell growth shown by the reduced colony formation in soft agar (right panel). (C) GSEA shows a negative enrichment of MYC signature genes, and a positive enrichment of neuronal genes after the silencing of *HAND2* in IMR32 cells for 72 hr. (D) Venn diagram shows that around 45% genes regulated by MYCN is also regulated by HAND2 in IMR32 cells. (E) GSEA shows a negative enrichment of MYC signature genes, and a positive enrichment of neuronal genes after knockdown of *HAND2* in LAN5 cells for 72 hr. (F) Venn diagram shows that around 33% genes regulated by MYCN is also regulated by HAND2 in LAN5 cells.

To investigate the long-term effect of losing *HAND2* in NB cells, we generated IMR32 CRISPR/Cas9 stable clones, with a control single guide RNA (sgCtrl) or a single guide RNA that targeted *HAND2* gene locus (*sgHAND2*) using a doxycycline (Dox) inducible CRISPR-Cas9 expression. Dox treatment decreased HAND2 protein levels in *sgHAND2* cells but not *sgCtrl* cells (Figure 2B, left panel) and this was accompanied by decreased cell proliferation (Figure S2F). Importantly, the Dox treatment resulted in reduced colony formation in soft agar in *sgHAND2* cells but not *sgCtrl* cells (Figure 2B, right panel), indicating a role of HAND2 in NB cell transformation.

Since HAND2 has not been systematically studied using genome-wide measurements and approaches, we investigated the HAND2 regulated genes by RNA-seq in control and *HAND2* silenced cells. The knockdown of *HAND2* resulted in a slight decrease (~20%) in MYCN protein (Figure S2G, left panel). Gene set enrichment analysis (GSEA) of the RNA-seq data showed that loss of *HAND2* resulted in a significant negative enrichment of MYC targets (Figure 2C, left panel) and a significantly positive enrichment of a neuronal gene signature (Figure 2C, middle and right panel). For comparison, we performed an RNA-seq experiment after knocking down *MYCN* in IMR32 cells for 72 hr. Silencing of *MYCN* for 72 hr increased HAND2 protein levels (Figure S2G, left panel). Consistent with the observation that loss of *HAND2* decreased the MYC gene signature, we found that around 44% of the genes regulated by MYCN were also regulated by HAND2 (Figure 2D, Table S1), suggesting a cooperative role between HAND2 and MYCN in regulating gene transcription. Similar results were observed in LAN5 cells when either *HAND2* or *MYCN* was silenced after the siRNA transfection (Figure S2G, right panel; Figure 2E, F, Table S2).

### Loss of *HAND2* affects regional epigenetic modification

To investigate how HAND2 regulates gene transcription at the epigenetic level, we silenced *HAND2* in IMR32 cells and performed ChIP-seq using HAND2, H3K27ac, H3K4me3, H3K27me3, RNA Pol II and RNA Pol II Ser2P antibodies. H3K27ac signal marks active enhancers, H3K4me3 and H3K27me3 mark the active, repressive, or poised chromatin states, and RNA Pol II Ser2P (S2) signal marks active transcriptional elongation. In parallel, we performed an ATAC-seq to evaluate the chromatin accessibility. Analysis of HAND2 ChIP-seq identified 25549 high-confidence HAND2 binding sites in the IMR32 cells that aligned with or without those transcriptional marks shown in the heatmaps (Figure 3A). Genomic Regions Enrichment of Annotations Tool (GREAT) (47) analysis of HAND2 binding sites showed that HAND2 predominantly bound in regions distal (>5 kb) from the transcription start site (TSS) (Figure 3B). Further peak distribution analysis showed that 2.3% of HAND2-binding sites were at the promoter regions (defined by ± 1kb of TSS), 46% of HAND2-binding sites were at the active enhancer regions (overlapped with H3K27ac peaks), and the remaining HAND2-binding sites were un-classified (silencer regions, weak enhancer, or poised enhancer regions) (Figure 3C). Homer *de novo* motif scanning of HAND2 peaks identified two highly enriched known HAND2 binding motifs (Figure 3D) identified in NB (22) or in human embryonic stem cell derived mesoderm cells (GSE61475). GREAT gene ontology (GO) biological process analysis of the HAND2 ChIP-seq data showed that the HAND2 peak-associated genes were enriched in sympathetic nervous system development and noradrenergic neuron differentiation (Figure 3E).

**Figure 3.**
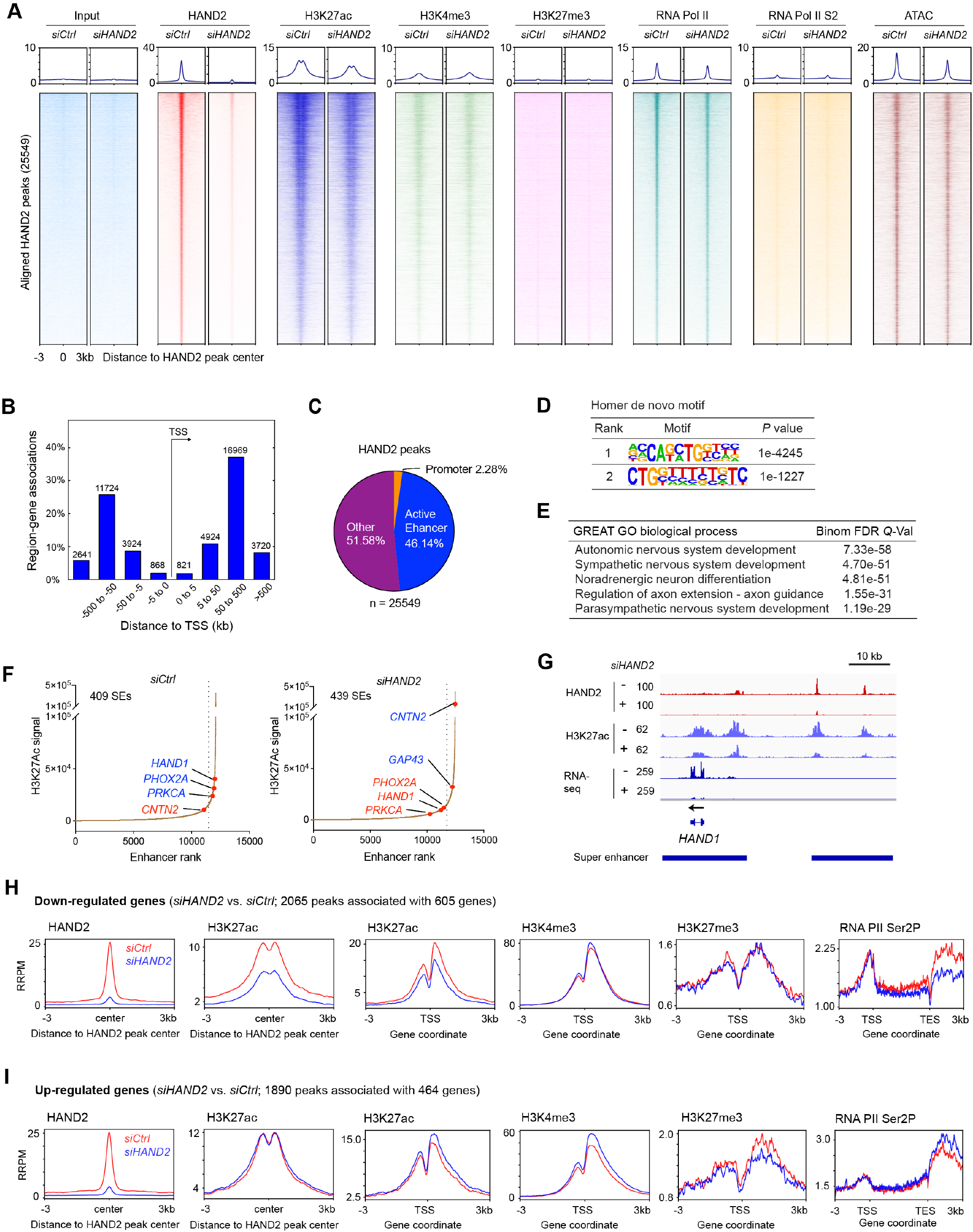
Loss of *HAND2* affects regional epigenetic modification. (A) Heatmaps of ChIP-seq data show the ranked HAND2 binding peaks (25,549) in siCtrl cells (narrow peaks, p<1e-7) and the aligned peaks of input, H3K27ac, H3K4me3, H3K27me3, RNA Pol II and ATAC-seq at HAND2 peak center before (siCtrl) and after (siHAND2) knocking down of HAND2 for 72 hr. Loss of *HAND2* results in a decrease of H3K27ac and RNA Pol II binding shown by the decreased signal of aligned peaks. (B) HAND2 mainly binds to genomic regions that are 5 kb up- or down-stream of TSS (transcription start site) by cis-regulatory regions prediction tool GREAT. (C) Venn diagram shows that around 46% of HAND2 binding sites are at active enhancer regions (overlapping with H3K27ac binding sites), 2.3% HNAD2 binding sites are at promoter regions (TSS ± 1 kb) and the other 50% unknown binding regions could be either poised enhancer or silencer regions. (D) Homer *de novo* motif scan shows the enrichment of two known HAND2 binding motifs. (E) HAND2 binding sites are associated with sympathetic nervous system development and neuron differentiation demonstrated by GREAT GO analysis (significant by region-based bionomial of 25549 peaks associated 18594 genes). (F) The silencing of *HAND2* affects super-enhancers establishment. (G) ChIP-seq signal tracks show that the knockdown of *HAND2* decreased signal of H3K27ac and RNA-seq at the *HAND1* gene locus. (H) Composite profile of ChIP-seq data shows a decreased average signal of H3K27ac at HAND2 peak center of genes down-regulated after the knockdown of HAND2 (left panel), a decreased signal of H3K27ac but not of H3K4m3 and H3K27me3 at TSS (middle panel), a decreased signal of RNA PolII Ser2P both at the gene body and in the region downstream of the poly-adenylation signal (TES, transcription end site). (I) Composite profile of ChIP-seq data shows no change of H3K27ac signal at HAND2 peak center of genes up-regulated after the knockdown of *HAND2* (left panel), but an increased signal of H3K27ac and H3K4m3, and a decreased signal of H3K27me3 at TSS (middle panel), an increased signal of RNA PolII Ser2P in the region downstream of the poly-adenylation signal (after TES), indicating active transcriptional elongation (right panel). RRPM: spike-in normalized, reference-adjusted reads per million mapped reads.

ChIP-seq and ATAC-seq heatmaps showed that majority of the ranked HAND2-centered peaks identified in siCtrl IMR32 cells aligned with the H3K27ac, RNA Pol II peaks and ATAC-seq peaks (Figure 3A), indicating most HAND2 binding regions are within accessible chromatin. No signals of H3K27me3 and weak signals of H3K4me3 were detected at the HAND2 peak center (Figure 3A), both of which were known to be restricted from enhancer regions. This observation was consistent with the majority of HAND2 peaks being at distal regulatory regions and active enhancer regions (Figure 3B,C). Knocking down of *HAND2* resulted in an almost complete loss of HAND2 signal that was accompanied by a decrease of H3K27ac and RNA Pol II signal at the HAND2 peak center regions (Figure 3A), suggesting that the loss of *HAND2* decreased enhancer activity. We also observed that the loss of *HAND2* resulted in a decrease of ATAC-seq signal, indicating a decrease in chromatin accessibility (Figure 3A), however, unlike a pioneer factor, the silencing of *HAND2* did not result in a complete loss of chromatin accessibility.

As a component of a CRC, HAND2 is driven by super-enhancers (SEs) and binds to the SEs of genes important for cell-identity (22,24). However, whether HAND2 plays in the establishment of SEs has not been investigated. SEs are large cluster of enhancers that bind high densities of transcriptional components and are marked by extensive stretches of strong signal of H3K27ac (48). We used the ROSE (rank ordering of SEs) analysis tool to assess how the loss of HAND2 affected SEs establishment. We identified 409 SEs (associated with 355 genes) in *siCtrl* and 439 SEs (associated with 382 genes) in *siHAND2* cells (Figure 3F). The loss of *HAND2* resulted in a loss of 87 SEs associated genes and an acquisition of 114 SEs associated genes with 267 SEs associated genes relatively unchanged (Table S3). Upon *HAND2* silencing, we identified a previously unexplored repressive role of CRC components in the establishment of SEs. Representative genes with lost SEs include *HAND1, PHOX2A* and *PRKCA* and representative genes with gained SEs include neuronal genes *GAP43* and *CNTN2* were found in the *HAND2* silenced cells (Figure 3F). Representative ChIP-seq signal tracks showed that the knockdown of *HAND2* resulted in a dramatic decrease of H3K27ac and RNA-seq signals at the *HAND1* gene locus (Figure 3G), and a dramatic increase of H3K27ac and RNA-seq signals at the *CNTN2* gene locus (Figure S3A).

To investigate how HAND2 regulates gene transcription, we focused on genes that had both HAND2 binding sites and were transcriptionally regulated by HAND2. In combination with HAND2 ChIP-seq results (in *siCtrl* cells) and RNA-seq results (*siHAND2* vs. *siCtrl*), we identified 605 genes directly down-regulated (associate with 2065 peaks) and 464 genes directly up-regulated (associate with 1890 peaks) after knocking down of *HAND2* (Table S4). By focusing on the directly “down-regulated genes” after the silencing of *HAND2*, the composite profile of ChIP-seq data at the HAND2 peak center showed that the average signal for H3K27ac was decreased when HAND2 was knocked down (Figure 3H); a decreased signal of H3K27ac but not of H3K4me3 and H3K27me3 was observed at TSS (Figure 3H) and a decreased signal for RNA PolII Ser2P was observed both within the gene body and in the region downstream of the poly-adenylation signal (Figure 3H), indicating a decrease of transcriptional elongation. Ingenuity pathway analysis of these “down-regulated genes” showed that the silencing of *HAND2* activated PTEN signaling and repressed cell cycle control of chromosomal replication signaling (Figure S3B). An evaluation of genes directly “up-regulated” after knocking down of *HAND2*, the composite profile of ChIP-seq data at the HAND2 peak center showed no change for H3K27ac signal (Figure 3I); an increased signal of H3K27ac and H3K4m3, and a decreased signal of H3K27me3 was observed at TSS (Figure 3I); an increased signal of RNA PolII Ser2P in the region downstream of the poly-adenylation signal was observed (Figure 3I). Ingenuity pathway analysis of these up-regulated genes after knocking down of HAND2 (HAND2 repressed genes) showed that these genes were associated with activated neuronal development related signaling (Figure S3C). Taken together, we found that HAND2 activates gene transcription mainly by increasing enhancer activity and transcriptional elongation, while HAND2 represses gene transcription mainly by decreasing the promoter activity and inhibiting transcriptional elongation.

### HAND2 facilitates MYCN enhancer invasion

To investigate the possible cooperation between HAND2 and MYCN on DNA binding, we first analyzed HAND2 and MYCN ChIP-seq data derived from IMR32 cells. We found that 70% of the MYCN binding sites that overlapped with HAND2 binding sites (MYCN common peaks) were at the active enhancer regions (overlapped with H3K27ac binding sites), while ~4% are at the promoter regions (Figure 4A). However, almost 50% of the MYCN binding sites that did not overlap with HAND2 binding sites (MYCN unique peaks) were at the promoter regions (Figure 4B). Homer *de novo* motif scan identified low-affinity E-box-like sequences (CANNTG) in the peaks MYCN shared with HAND2, while the MYCN unique peaks contain a high-affinity canonical E-box (CACGTG) (Figure 4A, B). This was consistent with the finding in tumor cells with high level expression of *MYC* or *MYCN* show binding to low-affinity E-box-like sequences within enhancers (11,12). GREAT GO analysis revealed that the genes associated with common MYCN and HAND2 sites are enriched in the regulation of sympathetic nervous system development and noradrenergic neuron differentiation, while the genes associated with unique MYCN peaks were enriched in RNA processing and ribosome biogenesis (Figure 4C, D). These results suggested a cooperative role of HAND2 and MYCN on the regulation of neuronal differentiation in NB cells.

**Figure 4.**
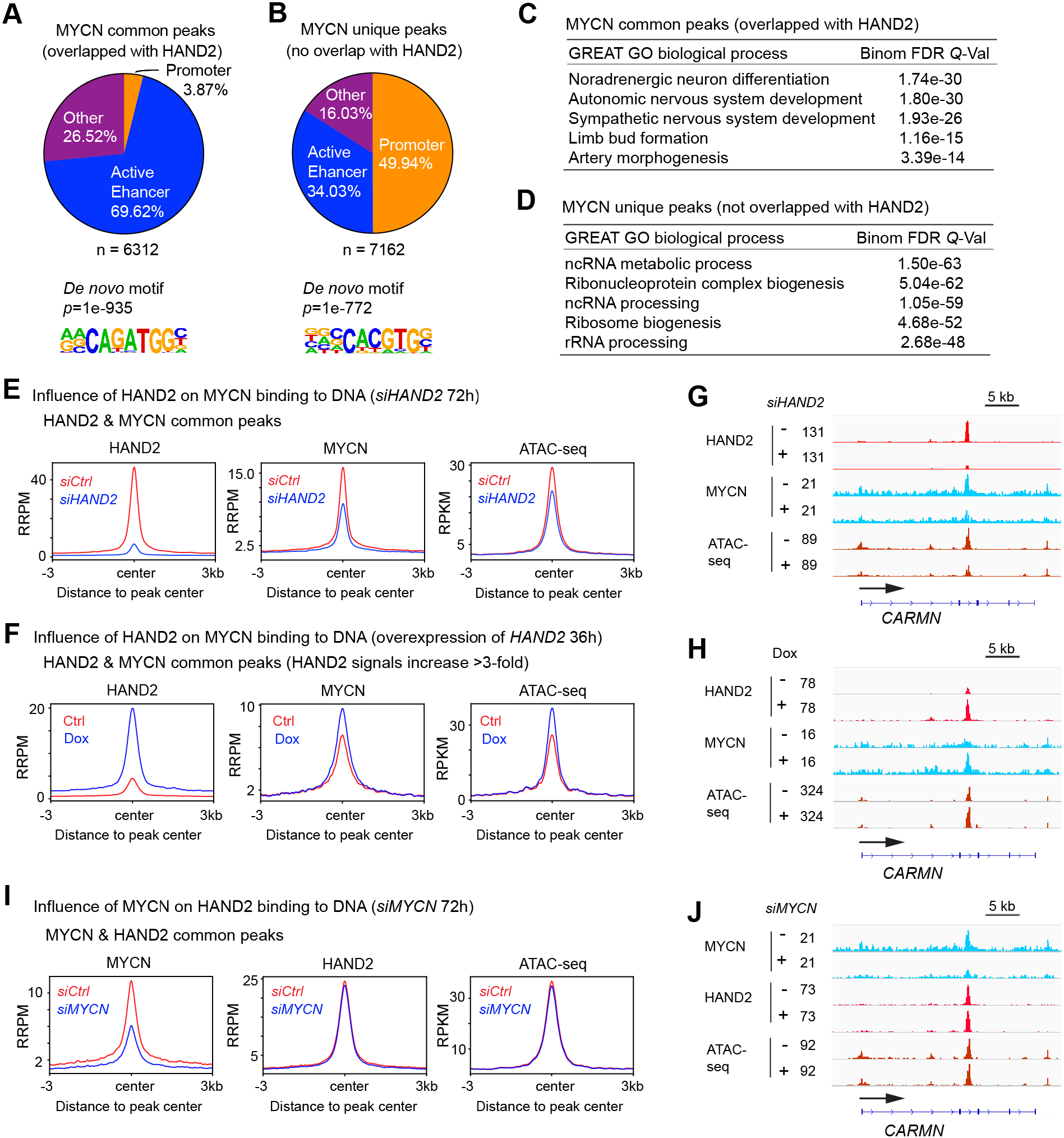
HAND2 assists MYCN to bind to DNA. (A) Venn diagram shows the distribution of MYCN binding sites that overlapped (>15 bp) with HAND2 binding sites (top panel). Homer *de novo* motif scan shows that the MYCN binding motif is a non-canonical E-box that known to be a weak MYCN binding motif. (B) Venn diagram shows the distribution of MYCN unique binding sites (no overlap with HAND2 binding sites). Motif scan shows that the MYCN binding motif is a canonical E-box that known to be a high-affinity MYCN binding motif. (C) GREAT GO biological process analysis shows that the MYCN and HAND2 overlapped peaks associated genes are enriched in noradrenergic neuron differentiation and sympathetic nervous system development. (D) GREAT GO biological process analysis shows that the MYCN binding peaks that are not overlapped with HAND2 associated genes are enriched in RNA processing. (E) Composite profile of ChIP-seq data shows a decreased average ChIP-seq signal of MYCN and ATAC-seq signal at the MYCN and HAND2 common sites after the knockdown of HAND2. (F) Composite profile of ChIP-seq data shows that the overexpression of HAND2 results in an increased average signal of MYCN at the MYCN and HAND2 common sites that accompanied by an increase of average ATAC-seq signal. Here only the HAND2 and MCYN common peaks with great than 3-fold increase of HAND2 ChIP-seq signals are included. (G) Signal tracks show that the knockdown of HAND2 results in a decrease of MYCN signal and ATAC-seq signal within the *CARMN* gene locus. (H) Signal tracks show that the overexpression of HAND2 results in an increased MYCN signal and ATAC-seq signal within the *CARMN* gene locus. (I) Composite profile of ChIP-seq data shows that the knockdown of MYCN has no effect on average HAND2 signal and the ATAC-seq signal at the MYCN and HAND2 common sites. (J) Signal tracks show that the knockdown of MYCN has no effect on HAND2 signal and ATAC-seq signal within the *CARMN* gene locus. RRPM: spike-in normalized, reference-adjusted reads per million mapped reads; RPKM: reads per kilobase per million mapped reads.

Next, we investigated the influence of HAND2 on MYCN binding to DNA using loss of function and gain of function studies in IMR 32 cells. When focused on the HAND2 and MYCN overlapping binding sites, there was a 40% decrease in the average ChIP-seq signal of MYCN (summit of the composite plot) when HAND2 was knocked down (Figure 4E). There was also a decrease in the average ATAC-seq signal at the HAND2 and MYCN common peaks after the silencing of *HAND2* (Figure 4E). After the knockdown of *HAND2*, the average MYCN ChIP-seq signal intensity decreased by 27.5% in the shared MYCN and HAND2 binding sites while the decrease was ~ 18.6% for the unique MYCN peaks (Figure S4A). The decrease of MYCN binding at the MYCN unique peaks during loss of HAND2 suggests a complex regulation of cooperative DNA binding between HAND2 and MYCN, which might be due to the nature of enhancer-enhancer and enhancer-promoter interactions. For gain of function studies of *HAND2*, we generated a stable Dox inducible, *HAND2* expressing IMR32 cell line (IMR32tetHAND2). After a 36 hr Dox treatment, we observed an up-regulation of HAND2 but not MYCN at the protein levels (Figure S4B). Consistent with the loss of function study, the gain of *HAND2* expression increased the average MYCN ChIP-seq signal at their common binding sites (Figure 4F) and this was accompanied by increased chromatin accessibility as indicated by an increase in the ATAC-seq signal (Figure 4F). Representative signal tracks within the *CARMN* gene locus showed that the knockdown of *HAND2* led to a decrease in the MYCN binding signal accompanied by a decrease in the ATAC-seq signal (Figure 4G), while the overexpression of *HAND2* led to an increase of MYCN binding signal and ATAC-seq signal (Figure 4H). The ATAC-seq results indicated that the overexpression of *HAND2* potentiates chromatin accessibility instead of de-compacting the completely closed chromatin.

To investigate whether the loss of *HAND2* affects MYCN binding at an earlier time point, we performed a ChIP-seq assay after knocking down *HAND2* for 24 hr. Western blot result showed that the knockdown of *HAND2* for 24 hr did not affect *MYCN* protein expression (Figure S4C). If one focused on all the HAND2 and MYCN overlapped binding sites, the loss of *HAND2* resulted in a 5% decrease of the summit of the average MYCN ChIP-seq signal at HAND2 peak center (Figure S4D, left panel). However, for the HAND2 and MYCN overlapping peaks with greater than a 4-fold decrease of HAND2 ChIP-seq signal, there was a 25% decrease of the summit of the average MYCN ChIP-seq signal (Figure S4D, middle panel), while the loss of *HAND2* for 24 hr did not affect MYCN ChIP-seq signal of those MYCN unique peaks that did not overlap with HAND2 binding sites (Figure S4D, right panel). When IMR32tetHAND2 cells were treated with Dox for 6 hr, HAND2 protein levels increased without an increase in MYCN protein levels (Figure S4E). Yet the overexpression of *HAND2* for only 6 hr resulted in an increase of the average MYCN ChIP-seq signal at HAND2 peak center with HAND2 signals increasing by 3 to 4-fold (Figure S4F). Representative signal tracks close to *TBX4* or *UQCRFS1* gene locus showed that the knockdown of *HAND2* caused a decrease of the MYCN ChIP-seq signal, while the overexpression of *HAND2* led to an increase of MYCN ChIP-seq signal in a time dependent manner on some but not all (indicated by different color boxes) the HAND2 & MYCN overlapping binding sites (Figure S4G, H).

Next, we investigated how loss of *MYCN* affects HAND2 binding to DNA. Composite profiles showed that the loss of *MYCN* did not affect average HAND2 ChIP-seq or ATAC-seq signal when focused on MYCN and HAND2 overlapping peaks (Figure 4I). Representative signal tracks within the *CARMN* gene locus showed that the knockdown of *MYCN* did not alter the HAND2 binding or ATAC-seq signal (Figure 4J). These results demonstrated that HAND2 assisted MYCN to bind to DNA, but MYCN did not alter HAND2 binding.

### CRC TF PHOX2B and GATA3 are not required for MYCN to bind to DNA

The cooperation between HAND2 and MYCN on DNA binding raised the question of whether other CRC TFs cooperatively bind to DNA. Since HAND2, PHOX2B and GATA3 have been broadly identified as CRC TFs in tested NB cell lines (22–24), we investigated whether HAND2 was required for GATA3 and PHOX2B to bind to DNA. The composite profile of ChIP-seq data at the HAND2 peak center showed a decreased average ChIP-seq signal of GATA3, which was accompanied by a decrease of ATAC-seq signal when *HAND2* was knocked down (Figure 5A, top panel), while the average signal of PHOX2B was not changed (Figure 5A, bottom panel). An example of such changes can be seen in the signal tracks close to the *LINC00682* gene locus (Figure 5B). The decrease of GATA3 ChIP-seq signal might be due to the decrease of total GATA3 protein levels caused by the silencing of *HAND2* (Figure S5A). Nevertheless, the knockdown of *HAND2* in IMR32 cells resulted in a decrease of global GATA3 DNA binding, which demonstrated a different type of cooperation between these two TFs on gene transcriptional regulation.

**Figure 5.**
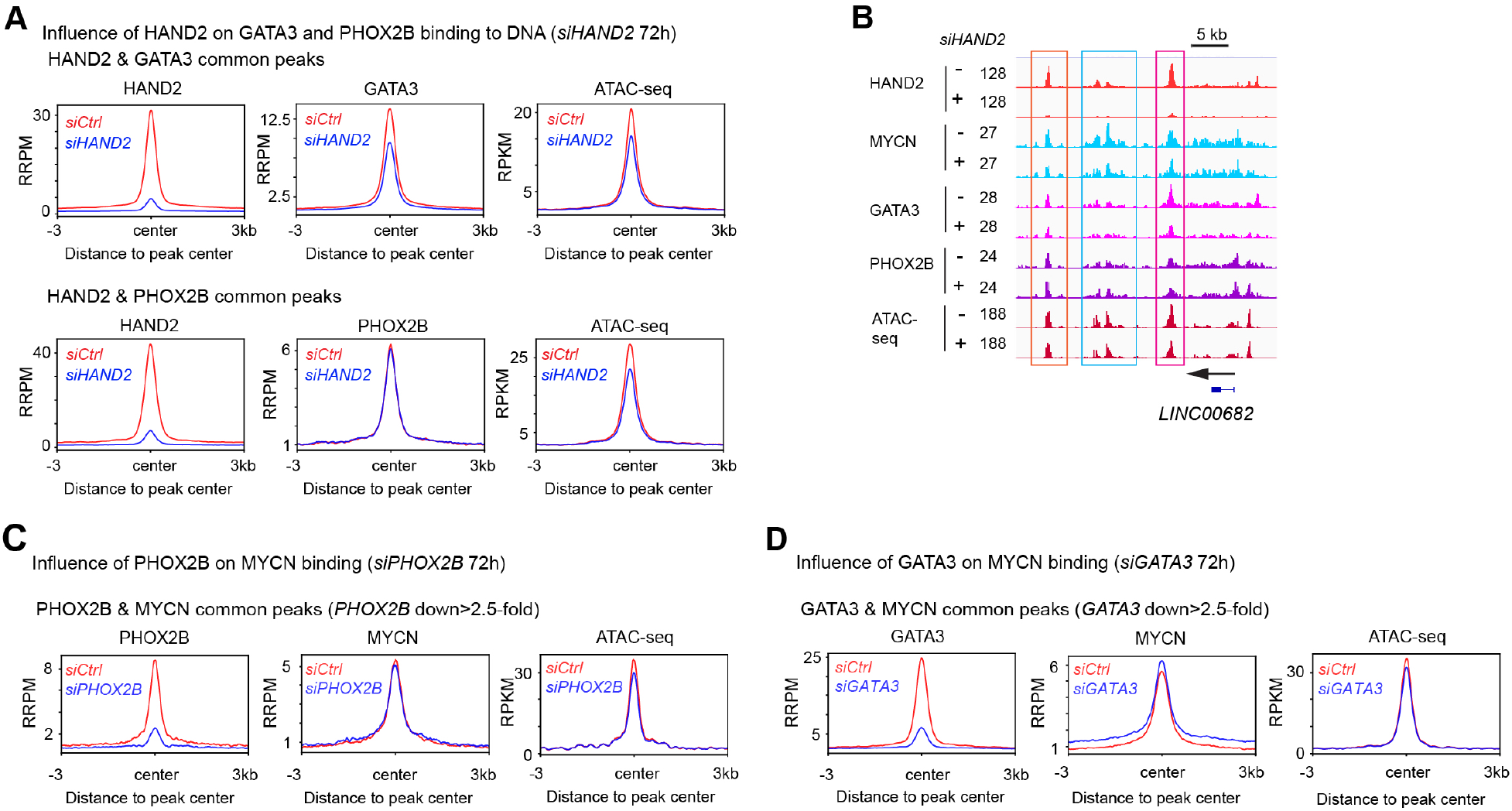
PHOX2B and GATA3 are not required for MYCN to bind to DNA. (A) Composite profile of ChIP-seq data shows that the knockdown of HAND2 results in a decreased average ChIP-seq signal of GATA3 and ATAC-seq signal at the HAND2 and GATA3 common binding sites (top panel) but has no effect on PHOX2B binding (bottom panel). (B) Signal tracks show that the knockdown of HAND2 results in a decrease of MYCN signal, GATA3 signal and ATAC-seq signal, but not PHOX2B signal next to the *LINC00682* gene locus. Peaks within the brown and cyan boxes indicate a decrease of ChIP-seq signals of HAND2, MYCN and GATA3, while peaks within the pink box indicate a decrease of HAND2 and GATA3 ChIP-seq signal. (C) Composite profile of ChIP-seq data shows almost no change of the average ChIP-seq signal of MYCN when focused on PHOX2B peaks with greater than 2.5-fold decrease of PHOX2B ChIP-seq signal after the silencing of *PHOX2B*, while a decrease of average ATAC-seq signal is observed at the PHOX2B and MYCN overlapped binding sites. (D) Composite profile of ChIP-seq data shows that the knockdown of GATA3 does not result in a decreased average signal of MYCN when focused on GATA3 peaks with greater than 2.5-fold decrease of GATA3 ChIP-seq signal after the silencing of *GATA3*.RRPM: spike-in normalized, reference-adjusted reads per million mapped reads; RPKM: reads per kilobase per million mapped reads.

Next, we knocked down *PHOX2B* or *GATA3* in IMR32 cells and performed ChIP-seq and ATAC-seq experiments to investigate whether these TFs were required for MYCN to bind to DNA. Western blot results showed that the knockdown of *PHOX2B* for 72 hr did not reduce MYCN protein levels while the knockdown of *GATA3* resulted in an increase of MYCN protein levels (Figure S5B). When focused on all the common peaks of PHOX2B and MYCN, the composite profile of ChIP-seq data at the PHOX2B peak center showed that the average signal of MYCN did not decrease (Figure S5C). Next, we tried to focus on those PHOX2B peaks with greater than 2.5-fold decrease of PHOX2B signal after the knockdown of *PHOX2B*. In this analysis, we observed that the average ChIP-seq signal of MYCN did not change, while a 10% decrease of the summit of the average ATAC-seq signal was observed (Figure 5C). Similar experiments and data analyses were done for *GATA3* gene. We found that the silencing of *GATA3* resulted in an increase of MYCN ChIP-seq signal at the GATA3 peak center when *GATA3* was knocked down (Figure 5D, Figure S5D), which was possibly due to the increase of total MYCN protein levels (Figure S5B). Taken together, our results are consistent with a model in which the CRC TF HAND2, but not PHOX2B or GATA3 is required for MYCN to bind to the distal regulatory DNA elements.

### HAND2 and MYCN cooperatively regulate chromatin accessibility

The “*indirect cooperative*” TF-DNA binding model proposes that the TFs recognize adjacent binding sites within regulatory regions, work together to compete with a nucleosome to access and cooperatively bind DNA, although no physical or direct protein-protein interaction occurs between these TFs (2,30,31). If HAND2 and MYCN cooperate in this way, the loss of both *HAND2* and *MYCN* should have a dramatic effect on nucleosome occupancy. To test this both *HAND2* and *MYCN* were knocked down simultaneously and the decrease in their protein expression was detected by western blot (Figure S6A). By focusing on the MYCN and HAND2 common binding sites, we stratified the ChIP-seq peaks into Group I, those that had a decreased MYCN ChIP-seq signal (>1.5-fold change); or Group II in which there was no change in the MYCN signal after the silencing of *HAND2*. Analysis of the resulting peaks (heatmap) showed that the loss of *HAND2* decreased MYCN binding and was accompanied by a decrease in the ATAC-seq signal in Group I regions (Figure 6A, left panel). In contrast after *HAND2* knockdown, the Group II regions were not associated with changes in MYCN binding or average ATAC-seq signal (shown in the heatmap, Figure 6A, left panel). The same results were also shown in composite plots (Figure 6A, right panel). One mechanism to explain these results is that when HAND2 binds to DNA, it facilitates the MYCN binding, then HAND2 and MYCN cooperate to compete with nucleosomes; whereas MYCN could not initiate this process, since the silencing of MYCN did not affect HAND2 binding (Fig. 4I) and the loss of *MYCN* alone did not affect chromatin accessibility within these regions (Figure 6A). Consistent with this hypothesis, the knockdown of both *HAND2* and *MYCN* led to greater decreases in chromatin accessibility shown by the decreased signal of ATAC-seq at the HAND2 and MYCN overlapped binding sites in both heatmaps and composite plots (Figure 6A). Next, we specifically focused on those HAND2 and MYCN common peaks in which there were no changes in MYCN binding and ATAC-seq signal after silencing *HAND2* alone (Figure 6A, Group II). In these Group II genomic regions, we found that the knockdown of either *HAND2* or *MYCN* alone had no effect on chromatin accessibility shown by the sustained ATAC-seq signal. It was only the loss of both *HAND2* and *MYCN* that decreased chromatin accessibility as detected by the decreased ATAC-seq signal (Figure 6A). This indicated that the co-existence of HAND2 and MYCN could compete with nucleosomes in these Group II genomic regions. Moreover, these results also suggest that for these Group II genomic regions, a third TF might play a role, thus, in the presence of this TF, the loss of either HAND2 or MYCN alone did not decrease the chromatin accessibility. When focused on all the active enhancer regions with ATAC-seq signal (ATAC-seq signal that overlapped with H3K27ac binding sites but not including TSS±1kb), the loss of both *HAND2* and *MYCN* resulted in a 50% decrease of the summit of the ATAC-seq signal (Figure S6B, left panel). Moreover, when focused on all the TSS regions, which were normally not co-bound by both HAND2 and MYCN, the loss of both *HAND2* and *MYCN* also resulted in a 50% decrease of the summit of the ATAC-seq signal (Figure S6B, right panel). This pointed to a global decrease in chromatin accessibility after loss of both HAND2 and MYCN, possibly through the disruption of enhancer-enhancer and enhancer-promoter interactions.

**Figure 6.**
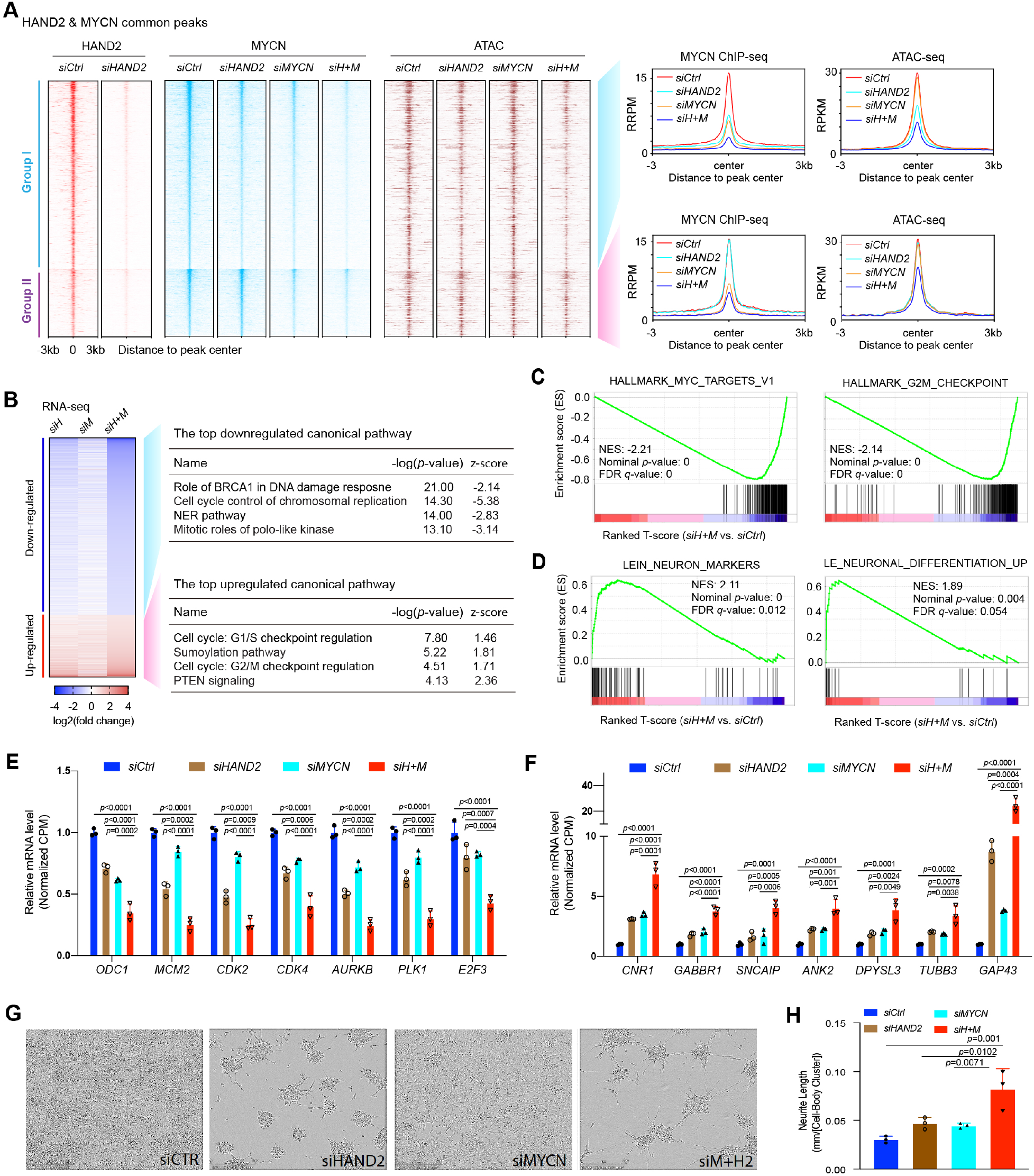
HAND2 and MYCN cooperatively regulate gene transcription. (A) Heatmaps (left panel) and composite profiles (right panel) of ChIP-seq and ATAC-seq results before and after the knockdown of *HAND2* (*siHAND2*), *MYCN* (*siMYCN*) or both (*siH+M*) in IMR32 cells (72 hr). These peaks are HAND2 and MYCN overlapped common peaks that grouped by either with a decreased MYCN ChIP-seq signal (>1.5-fold, 2629 peaks, Group I), or no change of MYCN ChIP-seq signal (<1.1-fold, 948 peaks, Group II) after the silencing of *HAND2*. In the heatmap, the peaks are ranked by MYCN intensity in *siCtrl*,and aligned by the HAND2 ChIP-seq peaks and ATAC-seq peaks. The decrease of ATAC-seq signal in both Group I and Group II could be observed when both *HAND2* and *MYCN* are silenced (*siH+M*). (B) Heatmap shows that the knockdown of both *HAND2* and *MYCN* using siRNAs (*siH+M*) results in greater effect on the down- or up-regulation of gene expression than knocking down each one alone (*siH* or *siM*) in IMR32 cells (72 hr), with more genes being down regulated (3964) and fewer genes being up-regulated (1381). Ingenuity canonical pathway analysis shows that the down regulated genes are enriched in the negative regulation of chromosomal replication and the up-regulated genes are enriched in the positive regulation of PTEN signaling. (C) GSEA shows that the knockdown of both *HAND2* and *MYCN* results in a negative enrichment of *MYC* target genes and genes involved in G2M checkpoint. (D) GSEA shows that the knockdown of both HAND2 and MYCN results in a positive enrichment of neuron markers and neuronal differentiation genes. (E) The silencing of both of *HAND2* and *MYCN* results in a more significant down-regulation of genes that required for G2M progression than the loss of each TF alone based on the RNA-seq results. The *p*-value indicated is calculated in one-way ANOVA. CPM: counts per million. (F) The silencing of both *HAND2* and *MYCN* leads to a more significant increase of neuronal differentiation genes than loss of each one alone. The *p*-value indicated is calculated in one-way ANOVA. (G) The knockdown of both *HAND2* and *MYCN* results in a significant increase of neurite length shown by the phase contrast image, and (H) the IncuCyte neurite-length assay. RRPM: spike-in normalized, reference-adjusted reads per million mapped reads; RPKM: reads per kilobase per million mapped reads.

### HAND2 and MYCN cooperatively regulate gene transcription

To determine the effect of loss of both *HAND2* and *MYCN* on global gene transcriptional regulation, we performed RNA-seq analysis. The silencing of both *HAND2* and *MYCN* in IMR32 cells led to greater changes in gene expression when compared with silencing of either *HAND2* or *MYCN* alone (Figure S6C, Figure 6B left panel, Table S5). Ingenuity pathway analysis showed that the knockdown of both *HAND2* and *MYCN* decreased transcription of the “mitotic signaling” pathway and activation of transcription in the “G2/M checkpoint regulation” and “PTEN signaling” pathways (Figure 6B right panel). GSEA showed that the combined knockdown resulted in a negative enrichment of *MYC* signature genes and G2M checkpoint signature genes (Figure 6C), and a positive enrichment of neuronal markers and neuronal differentiation genes (Figure 6D). For example, the silencing of both of *HAND2* and *MCYN* resulted in a more significant down-regulation of genes that are required for G2M progression and cell proliferation such as *CDK4, AURKB* and *PLK1* than loss of either TF alone (Figure 6E). Similarly, the silencing of both *HAND2* and *MYCN* led to a more significant increase in neuronal differentiation genes including *TUBB3, DPYSL3* and *GAP43* (Figure 6F). These results support the model that HAND2 cooperates with MYCN to regulate gene transcription. Consistent with the dramatic epigenome and transcriptome changes resulting from the silencing of both *HAND2* and *MYCN* expression, the loss of both of these TFs led to increased functional neuronal differentiation compared with loss of either one alone (Figure 6G,H).

### Targeting both MYCN and the CRC TFs is effective in suppressing NB tumor growth

CRC TFs are essential in NB cells (24). Our results demonstrated that MYCN and HAND2 cooperatively bind to DNA to regulate gene transcription and determine a malignant NB phenotype. These observations support the idea that the dysregulated TFs in cancer cells are relevant therapeutic targets (49–51). Thus, we investigated the effect of targeting both MYCN and CRC TFs in suppressing NB growth. As a proof of concept, here we selected the Aurora A kinase inhibitor (AURKAi) alisertib to target MYCN, and the HDAC inhibitor (HDACi) LBH589 to target CRC TFs. AURKA interacts with both MYCN and the SCF ubiquitin ligase to stabilize MYCN protein and AURKAi treatment increases degradation of MYCN in NB cells (52,53), however clinical trials of AURKAi showed limited efficacy against solid tumors (54). HDACi treatment is known to disrupt both super-enhancers (SEs) and CRCs, whereas the SEs often drive oncogenes in cancers and CRC components are found to be essential in cancers (24,48,55,56). However, repeated trials of HDACi alone has demonstrated limited success in treating solid tumors in clinical trials (57). Here we tested the combination of AURKAi and HDACi to treat NB cells and investigate the combination as a potential therapeutic strategy.

We found that the treatment of NB cells with alisertib and LBH589 was more effective in reducing the viable cell number in *MYCN*-amplified cell lines compared to *MYCN* single-copy NB cell lines that have low levels of both MYCN and c-MYC (Fig. 7A, Fig. S7A), as only 0-50% viable *MYCN*-amplified cells were observed after 10 nM alisertib or 10 nM LBH589 treatment, while over 70%*MYCN* single-copy cells were still alive after the same dose of drug treatment (Fig. 7A). Half-maximal inhibitory concentration (IC50) assays clearly showed that *MYCN*-amplified cell lines were more sensitive to the LBH589, with IC50 values averaging 4x lower than those found in *MYCN* single-copy cell lines (8.5 nM vs. 33.5 nM, Fig. S7B, left panel). Moreover, *MYCN*-amplified cell lines were far more sensitive to alisertib compared to *MYCN* single-copy cell lines, since the average IC50 values of *MYCN*-amplified cells was 7.8 nM, while there were still more than 50% viable *MYCN* single-copy cells existed even after 1000 nM alisertib treatment for 72 hr (Fig. S7B, right panel). These results indicated potent and selective effects of both HDACi and AUKRAi in *MYCN*-dysregulated cells. Importantly, the combination treatment using alisertib and LBH589 synergistically reduced viable NB cell number with average bliss synergy scores greater than 10 across a range of doses in IMR32, IMR5 or KCNR cells (Fig. 7B). IncuCyte cell confluence assays confirmed the synergistic effects of alisertib + LBH589 treatment on cell proliferation in *MYCN*-amplified NB cell lines (Fig. 7C). As expected, 6 hr of HDACi LBH589 treatment resulted in a decrease of the SE driven CRC components *HAND2*, *PHOX2B* and *GATA3* at the mRNA levels in all the tested NB cell lines (Fig. S7C). The decrease of SE driven *MYCN* expression at mRNA levels was observed in IMR32 cell line after 6 hr LBH589 treatment (Fig. S7C). As previously reported the inhibition of AURKA destabilized MYCN (52,53) and we found that the 48 hr alisertib treatment resulted in a 30-50% decrease of MYCN at protein levels in NB cells compared to the MYCN protein levels in the controls (Fig. 7D). LBH589 treatment resulted in a decrease of both MYCN and CRC TFs protein levels compared with control, while the effect was TF and cell line dependent, and decreases only approaching 50% in most of cell lines evaluated (Fig. 7D). These results indicated that AURKAi or HDACi alone was not very effective at hitting these targets. However, the combination of alisertib and LBH589 treatment caused ~80% decrease of MYCN and the CRC TFs at the protein levels after 48 hr treatment in all the tested cell lines (Figure 7D), indicating that this combined treatment was far more potent at inhibiting these targets than single agent treatment. Of note, unlike the specific genetic silencing of HAND2 and MYCN, we did not observe a significant increase of neuronal differentiation, which might be due to the broad effect of alisertib + LBH589 may have on targeting many super-enhancers that drive essential NB genes, which resulted in a dominant effect on cell proliferation and/or cell survival.

**Figure 7.**
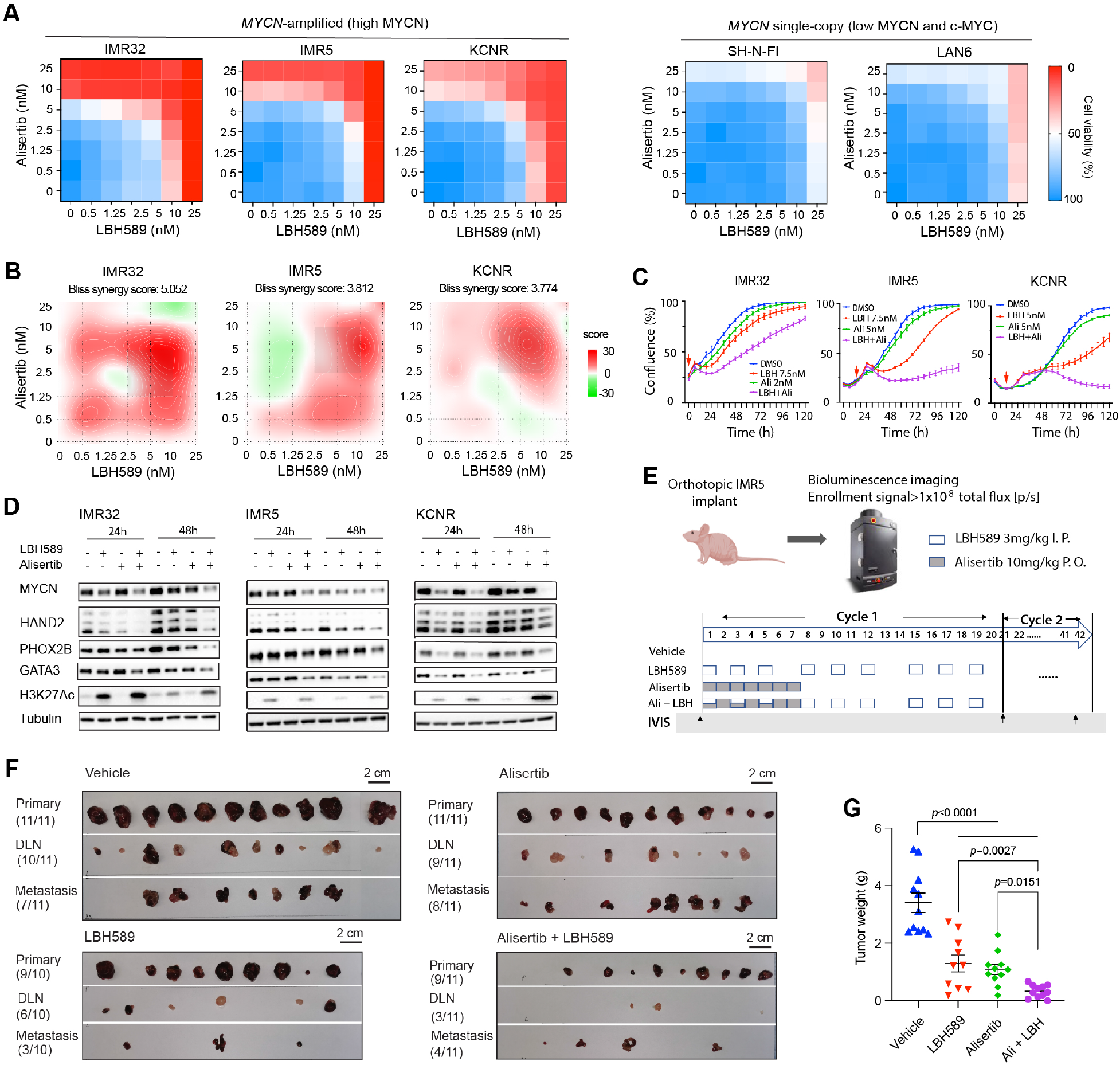
Targeting both MYCN and CRC TFs. (A) Heatmaps show the percentage of cell viability after different dose of HDAC inhibitor LBH589 and Aurora A kinase inhibitor alisertib treatment in *MYCN*-amplified NB cell lines and *MYCN* single-copy cell lines. Cells are treated with the drugs for 72 hr and cell viability is measured by CellTiter-Glo Cell Viability Assay. (B) SynergyFinder online tool is used for bliss synergistic analysis to evaluate the synergistic effect of the combination treatment in *MYCN*-amplified cell lines shown in (A). (C) IncuCyte cell confluence assays show the synergistic effect of the alisertib (Ali) + LBH589 (LBH) treatment on cell proliferation (% confluency) over time. The red arrow is the time point of adding compounds. (D) Western blot analysis shows the protein levels of MYCN and CRC TFs in IMR32, IMR5 and KCNR cells that treated with LBH589 (LBH), alisertib (Ali) alone or in combination for 24 hr and 48 hr. The dose of the drugs used here is the same as shown in (C). (E) Schematic diagram to show the strategy of drug treatment in orthotopic IMR5-GFP-Luc implanted xenografts. Cycle 1 is from day 1 to day 21, and cycle 2 is from day 22 to day 42, which is a repeat of drug treatment of cycle 1. The two cycles are indicated by black arrows. IVIS: *in vivo* imaging system. (F) Images of the tumors collected from each of the mice treated with vehicle or different drugs. DLN: draining lymph nodes. (G) Tumor weight measurement shows a significant decrease of the tumor weight of the drug treatment groups compared to the tumor weight of the vehicle treatment group. The tumor weight of the alisertib + LBH589 treatment group is significantly lower than the tumor weight of the alisertib or LBH589 single drug treatment group, while the tumor weight of the vehicle treatment group is significantly higher than the tumor weight of the rest groups. The *p*-value indicated is calculated in one-way ANOVA.

Next, we evaluated the activity of alisertib combined with LBH589 in preclinical xenograft models with established orthotopic IMR5-GFP-Luc. Animals were randomized according to their bioluminescence signal and received therapy with two treatment cycles of the following groups: Vehicle; LBH589 (LBH) alone; alisertib (Ali) alone; and alisertib and LBH589 (Ali + LBH) in combination (Fig. 7E). A small cohort of mice of each treatment arm were used for pharmacodynamics (PD) study and tumors were collected from these mice after 28 hr of the first dose of drug (see the method section for detailed information). Protein levels of MYCN and CRC components in collected tumors were heterogeneous among the mice in each group while the overall levels of these proteins were lower in the drug treatment groups compared with the vehicle treatment group (Fig. S7D). Moreover, increased H3K27ac signal was observed in the groups of mice treated with either LBH589 alone or LBH589 + alisertib (Fig. S7D). These results demonstrated that the doses of drugs used were sufficient to hit the intended targets *in vivo*. Animals across all groups did not exhibit any signs of overt toxicity (Fig. S7E). Importantly, tumor size (Fig. 7F) and tumor weight (Fig. 7G) measurements revealed that the combination of alisertib and LBH589 treatment was superior to either single agent alone at suppressing tumor growth. Finally, fewer tumors were detected in the draining lymph nodes (DLN) and metastatic sites (e.g., liver, intra-abdominal sites) in the combination alisertib + LBH589 arm (31.8%) of the study when compared to alisertib alone arm (77.3%) or LBH589 alone arm (45%) (Fig. 7F). Our results demonstrated that the combination therapy of alisertib + LBH589 was effective in *MYCN*-amplified NB.

## DISCUSSION

Combinatorial control of gene expression by a small group of TFs is critical to establish and maintain the cancer cell identity (2,5,58). In this study, we identify that the CRC TF HAND2 assists MYCN in enhancer binding, and that these two TFs cooperatively regulate the global gene expression that is required to maintain the malignant state of NB cells. Loss of both *HAND2* and *MYCN* dramatically decreases chromatin accessibility, indicating that the presence of both HAND2 and MYCN competes with nucleosomes. Importantly, a combination of AURKAi and HDACi treatment of NB cells more potently reduces the protein levels of both MYCN and CRC TFs than the treatment of NB cells with each drug alone, which results in a more effective inhibition of NB tumor growth. This result indicates that the combination of a precision therapy of targeting MYCN with epigenetic therapy that targets SEs-driven TFs is a promising strategy for *MYCN*-amplified NB therapy.

Our study shows the selective influence of HAND2 on MYCN-enhancer binding as the loss of *HAND2* results in a decrease of genome bound MYCN while the overexpression of *HAND2* leads to an increase of genome bound MYCN. Zeid et al 2018 identified that MYCN invades enhancers but the mechanism by which MYCN was able to invade enhancers was not identified (12). NB arise from dysregulation during sympathoadrenal cell lineage development in which PHOX2A, PHOX2B, GATA3, HAND1 and HAND2 interact as a network to specify cellular identity (17,21). HAND2, PHOX2B and GATA3 are highly expressed in NB and form a CRC to cooperate with MYCN to determine a noradrenergic NB phenotype (24). Here we find that although both HAND2, PHOX2B and GATA3 interact with MYCN, only the loss of HAND2 decreases MYCN enhancer binding. In addition to assist MYCN to bind to DNA, other studies showed that HAND2 increases PHOX2A binding affinity on the DBH promoter and synergistically activates a DBH promoter-reporter in the noradrenergic cell type (16,59). This suggests a general role HAND2 plays in facilitating TF-DNA binding. Importantly, the silencing of both *HAND2* and *MYCN* simultaneously in NB cells results in greater changes in gene expression when compared with the silencing of either *HAND2* or *MYCN* alone. These results demonstrate that HAND2 facilitates MYCN invasion into enhancers to cooperatively regulate global gene expression.

Our evidence indicates that HAND2 facilitates MYCN enhancer invasion via an “*indirect cooperative*” TFs-DNA binding model. The Co-IP and western blot analyses indicate MYCN interactions with HAND2 require nucleic acids diminishing the likelihood of a “*direct cooperative*” TFs-DNA binding model, which depends on direct protein-protein interaction (1,2,27–29). In the “*indirect cooperative*” TFs-DNA binding model, TFs recognize adjacent binding sites within regulatory regions to compete with nucleosomes to gain access and bind DNA simultaneously or sequentially (2,30–33). Our ATAC-seq results show that the loss of *HAND2* partially decreases chromatin accessibility at the HAND2 binding sites, whereas the silencing of both of *HAND2* and *MYCN* results in a more dramatic decrease of chromatin accessibility than the silencing of either TF alone. Moreover, the overexpression of *HAND2* in IMR32 cells increases the chromatin accessibility as shown by the increased ATAC-seq signal at the HAND2 and MYCN overlapped binding sites. These results indicate that HAND2 and MYCN cooperate to compete with nucleosomes. Pioneer TFs can bind to nucleosomal DNA within closed chromatin, open the chromatin and facilitate the subsequent binding of additional TFs (60,61). However, our results do not support a pioneer TF role for HAND2 as typically the loss of pioneer factors such as Oct4 or Pax7 alone, leads to a complete loss of chromatin availability at the majority of their respective binding sites (62,63). Taken together, we identify that HAND2 and MYCN achieve specific DNA binding through an “*indirect cooperative*” TF-DNA binding mechanism. Ultimately, a future crystal structure of HAND2/MYCN/DNA ternary complex would be beneficial in defining the molecular interactions between HAND2 and MYCN on DNA binding.

One of the elements missing from our understanding of the CRC networks in gene transcriptional regulation is an understanding of the dynamics of TF-DNA binding needed to implement the transcriptional circuits. A CRC is formed by TFs that collaborate to form a feed-forward interconnected regulatory loops and co-occupy regulatory regions of their target genes to establish and maintain cell identity (40,64). The finding that the CRC component HAND2 assists MYCN to bind to DNA adds another layer to our understanding of how CRC TFs form a combinatorial transcriptional regulatory circuit to determine a cell identity in health and diseases. Efforts to re-wire oncogenic circuits require an understanding of the TF-DNA binding interactions. Unlike previous studies that used *in vitro* cell free models or focused on a few genes when used *in vivo* cell models, we show at a genome wide level the existence of “*cooperative*” TF-DNA binding. Through this mechanism, HAND2 assists MYCN to bind DNA and cooperatively regulate global gene expression. This finding provides a rational for targeting multiple TFs simultaneously to increase the disruption of this “combinatorial transcription regulatory network” in cancer cells for enhanced therapeutic efficacy. Indeed, our combination of AURKAi and HDACi designed to target both MYCN and CRC TFs could efficiently suppress *MYCN*-amplified NB growth and has important therapeutic implications for this devastating pediatric tumor.

## DATA AVAILABILITY

All the home generated RNA-seq, ChIP-seq and ATAC-seq datasets can be found in the Gene Expression Omnibus (GEO) database. GEO accession number for data generated in this study is GSE183641, and the subseries that are linked to GSE183641 are GSE183609 and GSE183636 for RNA-seq, GSE184057, GSE184058 and GSE184059 for ChIP-seq, GSE184229 and GSE184232 for ATAC-seq. GEO accession number for publicly available ChIP-seq data derived from BE2C cells is GSE94822.

## SUPPLEMENTARY DATA

Supplementary data include one single PDF file that includes 7 supplementary figures and 8 supplementary tables in MS Excel.

## ACKNOWLEDGEMENTS

We thank Bao Tran, Jyoti Shetty and Yongmei Zhao from NCI Sequencing Facility for DNA and RNA sequencing. This work utilized the computational resources of the NIH HPC Biowulf cluster (http://hpc.nih.gov). The graphical abstract was created with BioRender.com.

## FUNDING

This work was funded by the Center for Cancer Research, Intramural Research Program at the National Cancer Institute.

## CONFLICT OF INTEREST

The authors declare no competing interests.

## TABLE AND FIGURES LEGENDS

**Table S1**. Genes regulated by HAND2 or MYCN in IMR32 cells.

**Table S2**. Genes regulated by HAND2 or MYCN in LAN5 cells.

**Table S3**. Super-enhancers associated genes identified in *siCtrl* cells and *siHAND2* cells.

**Table S4**. HAND2 directly activated and repressed genes.

**Table S5**. Genes altered after silencing both *HAND2* and *MYCN*.

**Table S6**. Primer sequences used for realtime PCR.

**Table S7**. Sequence information of the *siRNAs* and *sgRNAs*.

**Table S8**. List of antibodies used in western blot, co-IP and ChIP-seq.

## Notes

### Competing Interest Statement

The authors have declared no competing interest.

