## Supplementary Figures for "A “*cooperative collaboration*” between HAND2 and MYCN shapes neuroblastoma cell identity"

**Running title: It takes a HAND(2) for MYCN to invade enhancers**

Man Xu^1^, Ming Sun^1^, Xiyuan Zhang^1^, Rosa Nguyen^1^, Haiyan Lei^1^, Jack F. Shern^1^, Carol J. Thiele^1*^ and Zhihui Liu^1*^

^1^Pediatric Oncology Branch, National Cancer Institute, Bethesda, MD, USA

^*^To whom correspondence should be addressed. Tel: 1-240-858-3857; Fax: 1-301-451-7052;. Correspondence may also be addressed to Carol J. Thiele. Tel: 1-240-858-3849; Fax: 301-451-7052;

**Supplementary Figures**

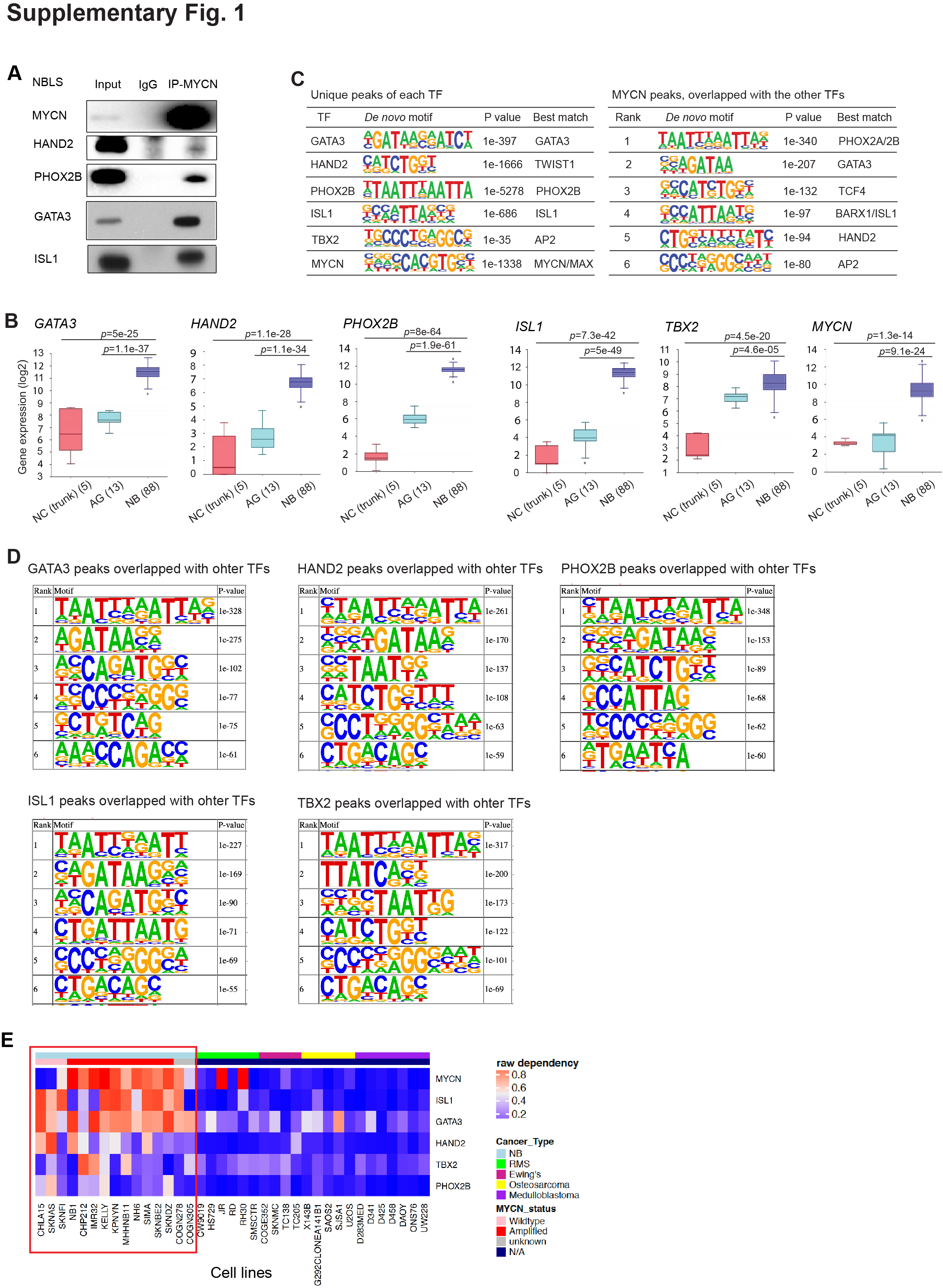

**Figure S1 (Extension of Figure 1). Association between NB CRC components and MYCN.** (A) MYCN interacts with CRC TFs in NBLS cells (single *MYCN* copy) shown by co-IP of MYCN protein followed by western blot. (B) The mRNA levels of CRC components and *MYCN* in human normal neural crest (NC), adrenal gland (AG) and neuroblastoma tumors (Versteeg cohort) are analyzed by the R2 database. Data are presented as box and whisker plots with middle lines indicating medians and whiskers representing the 25^th^ and 75^th^ percentiles. The *p*-value indicated is calculated in one-way ANOVA. The graph is generated by R2 database. (C) HOMER *de novo* motif scan of unique peaks of each CRC components and MYCN (left panel), as well as MYCN peaks that overlapped with the rest of the CRC components (right panel). (D) HOMER *de novo* motif scan of the binding sites of each component of the CRC that overlapped with the rest of the CRC components and MYCN. The discovered top ranked motifs are enriched in PHOX2A/PHOX2B and GATA3 binding motif and non-classical E-box in addition to their own specific binding motif, which indicates that these motifs are adjacent to each other and enriched at these regions. (E) Heatmap shows the dependency of different types of pediatric cancer cell lines on the CRC components and MYCN. NB cell lines (within the red box) are most preferentially dependent on these TFs when compared to other cancer types. Data mined from Project Achilles genome wide CRISPR-Cas9 screen.

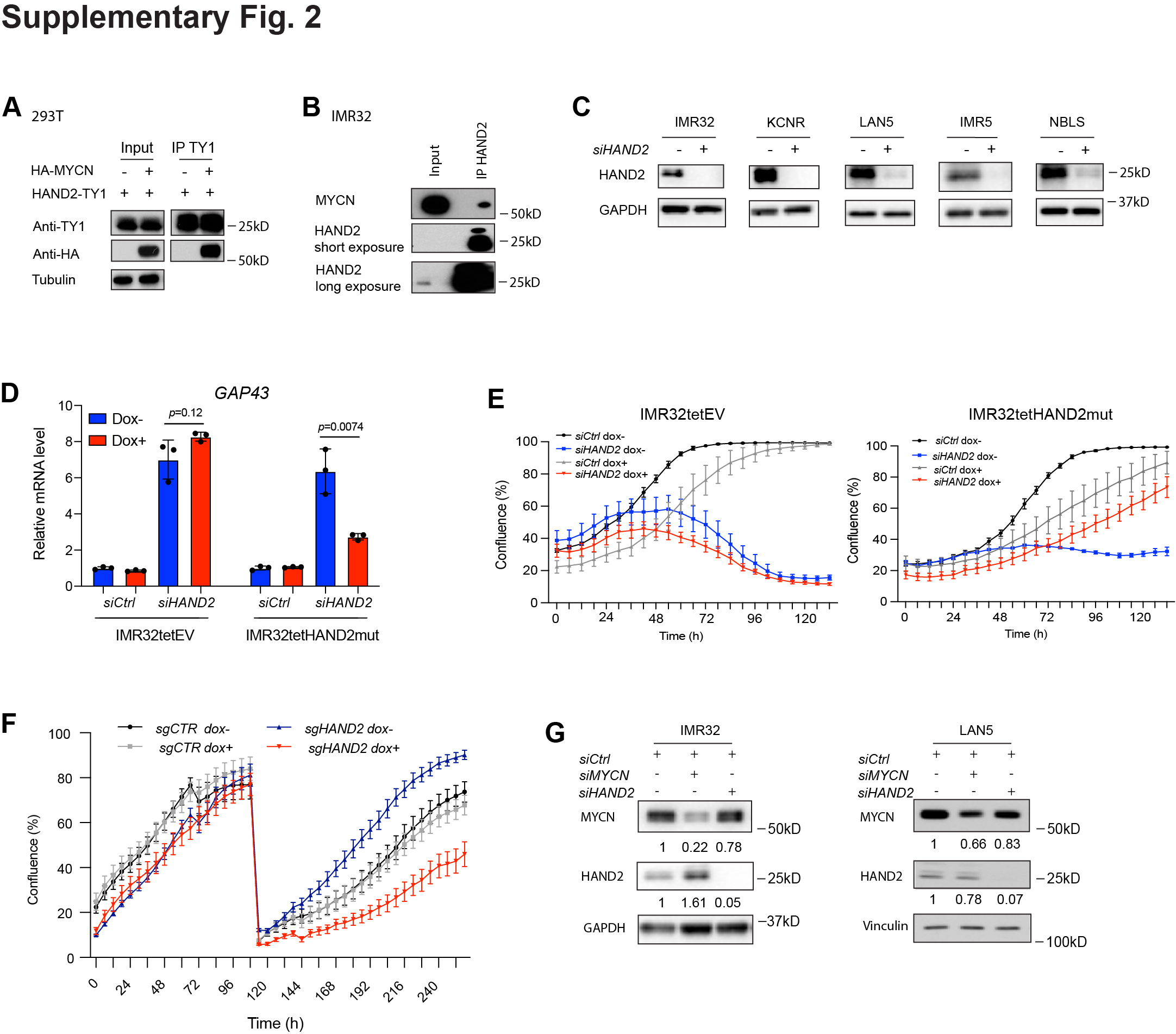

**Figure S2 (Extension of Figure 2). CRC component *HAND2* regulates the expression of *MYCN* targets and is essential in NB.** (A) Co-IP of HAND2 protein results in the pulldown of MYCN when they were transiently overexpressed in 293T cells shown by western blot assay. (B) Co-IP of endogenous HAND2 protein results in the pulldown of endogenous MYCN detected by western blot assay. (C) The knockdown of *HAND2* in different NB cell lines results in a decrease of HAND2 expression at protein levels shown by western blot assay. (D) Realtime PCR shows that the knockdown of *HAND2* results an increase of *GAP43* mRNA levels (increased by ~6.4-fold in both IMRtet32EV cells and IMR32tetHAND2mut cells without Dox treatment), while this up-regulation of *GAP43* mRNA levels in IMRtet32HAND2mut cells is attenuated when siRNA resistant HAND2 mutant is induced (Dox+) (increased by ~2.7-fold). The *p*-value indicated is calculated in Student’s t-test. (E) IncuCyte cell confluence assay shows that the suppression of cell proliferation after knocking down *HAND2* (siHAND2) could be rescued when siRNA resistant HAND2 mutant was induced by Dox. Note: Dox toxicity was observed in both IMR32tetEV and IMR32tetHAND2mut cells since Dox was added to the cells immediately after the electroporation. (F) Depletion of *HAND2* through Dox inducible Crispr/Cas9 and single-guide RNA (sgRNA) that targets *HAND2* but not control (sgCtrl) results in a decrease of cell proliferation based on cell confluence assay. Different group of cells with were passaged and re-plated at the same ratio when they reach 80% confluency. (G) The knockdown of *HAND2* and *MYCN* in IMR32 or LAN5 cells results in a decrease of their expression at protein levels shown by western blot assay.

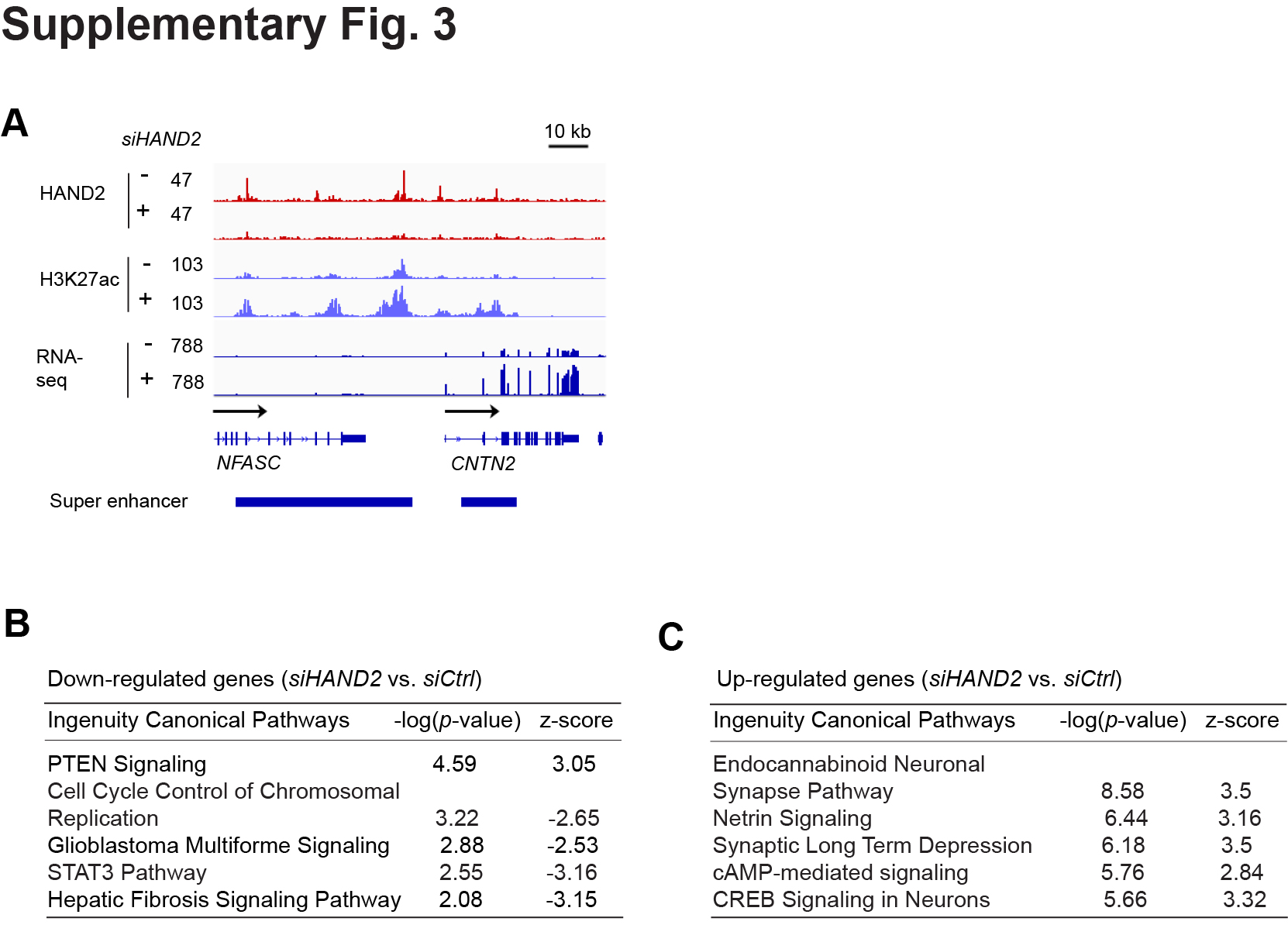

**Figure S3 (Extension of Figure 3). Loss of *HAND2* affects regional epigenetic modification.** (A) ChIP-seq signal tracks show that the knockdown of *HAND2* decreased signal of H3K27ac and RNA-seq around the *CNTN2* gene locus. (B) The knockdown of *HAND2* leads to the positive enrichment of PTEN signaling and negative enrichment of cell cycle control of chromosomal replication signaling demonstrated by ingenuity canonical pathways analysis. (C) The knockdown of *HAND2* leads to the positive enrichment of neuronal development related signaling demonstrated by ingenuity canonical pathways analysis.

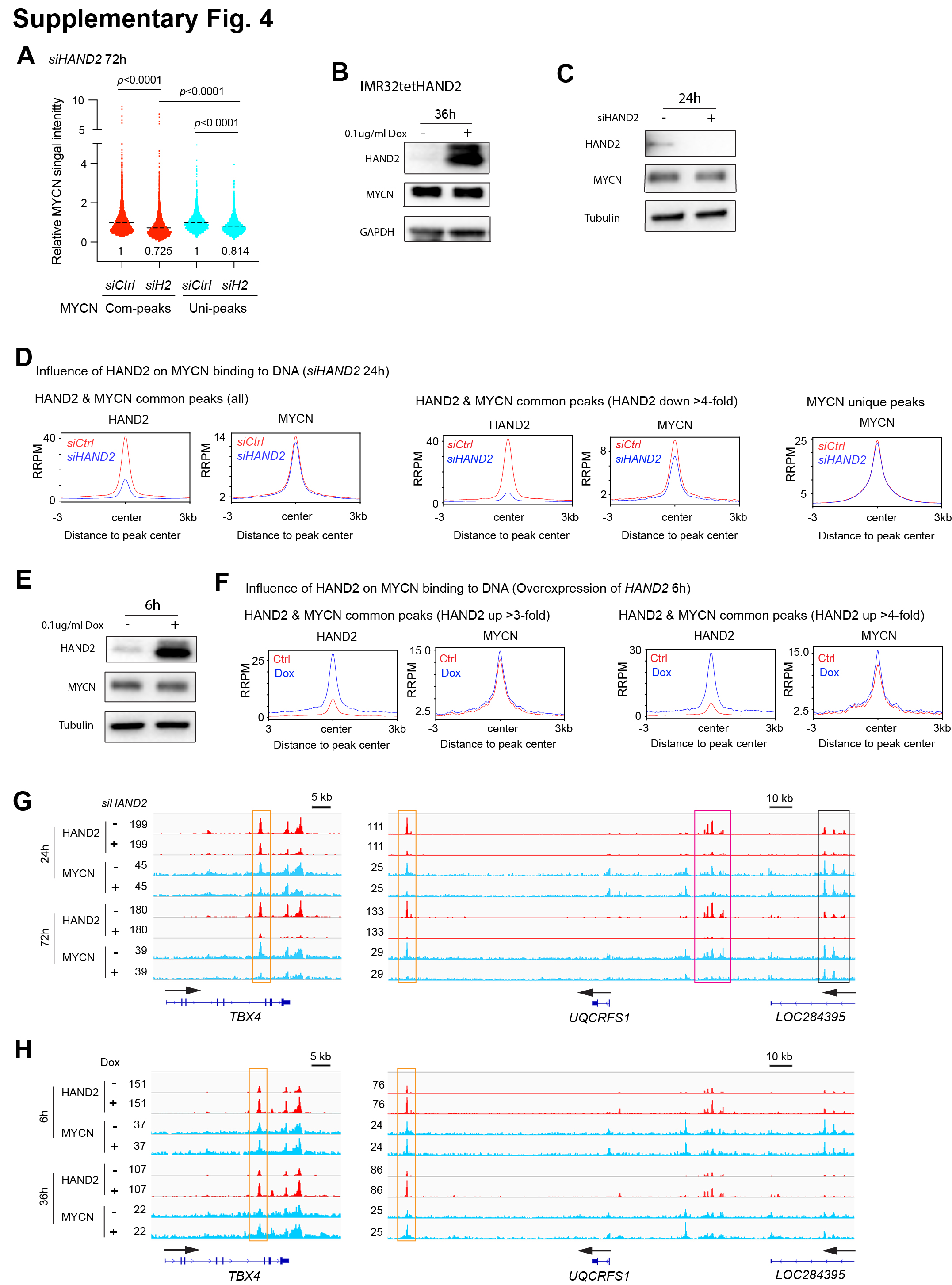

**Figure S4 (Extension of Figure 4). HAND2 assists MYCN to bind to DNA.** (A) Relative MYCN ChIP-seq signal intensity changes within the MYCN and HAND2 overlapped binding sites (Com-peaks), as well as the MYCN unique binding sites (Uni-peaks) after silencing of *HAND2* in IMR32 cells. The decrease of the MYCN signal within the MYCN & HAND2 common peaks is significantly greater than the decrease of the MYCN signal within the MYCN unique peaks. Note: Com-peaks represent MYCN and HAND2 overlapped peaks, Uni-peaks represent MYCN binding sites that not overlapped with HAND2 binding sites; the dot line indicates relative mean signal intensity, and the numbers are labeled under each of the violin plot. The *p*-value indicated is calculated in Student’s t-Test. (B) The overexpression of HAND2 in IMR32 cells for 36 hr results in an increase of HAND2 at protein levels without affecting MYCN protein levels shown by western blot assay. (C) The knockdown of HAND2 in IMR32 cells for 24 hr results in a decrease of HAND2 at protein levels without affecting MYCN protein levels shown by western blot assay. (D) Composite profile of ChIP-seq data shows the changes of average HAND2 and MYCN signals after the silencing of *HAND2* when focused on all the HAND2 and MYCN common binding sites (left panel); when focused on the HAND2 and MYCN common binding sites with >4-fold decrease of HAND2 signal, the silencing of *HAND2* results in a decrease of MYCN signal (middle panel); while the silencing of *HAND2* does not affect MYCN signal when focused on MYCN unique binding sites (MYCN peaks that not overlapped with HAND2 peaks). (E) The overexpression of *HAND2* in IMR32 cells for 6 hr results in an increase of HAND2 at protein levels without affecting MYCN protein levels shown by western blot assay. (F) Composite profile of ChIP-seq data shows that the overexpression of *HAND2* for 6 hr results in an increase of average MYCN signal when focused on the HAND2 and MYCN common binding sites with great than 3-fold (left panel) or 4-fold (right panel) increase of HAND2 signal. (G) Signal tracks show that the knockdown of *HAND2* for 24 hr or 72 hr results in a decrease of MYCN signal next to the *TBX4* gene locus (left panel), as well as the *UQCRFS1* and *LOC284395* gene locus (right panel). Peaks within the orange box and pink box indicate a decrease of MYCN ChIP-seq signal after the silencing of *HAND2* for both 24 hr and 72 hr, while peaks within the black box indicate a decrease of MYCN ChIP-seq signal only after 72 hr silencing of *HAND2*. (H) Signal tracks show that the overexpression of HAND2 for 6 hr or 36 hr results in an increase of MYCN signal next to the *TBX4* gene locus (left panel), as well as the *UQCRFS1* and *LOC284395* gene locus (right panel). Peaks within the orange box indicate an increase of MYCN ChIP-seq signal after the overexpression of *HAND2* for both 6 hr and 36 hr. RRPM: spike-in normalized, reference-adjusted reads per million mapped reads.

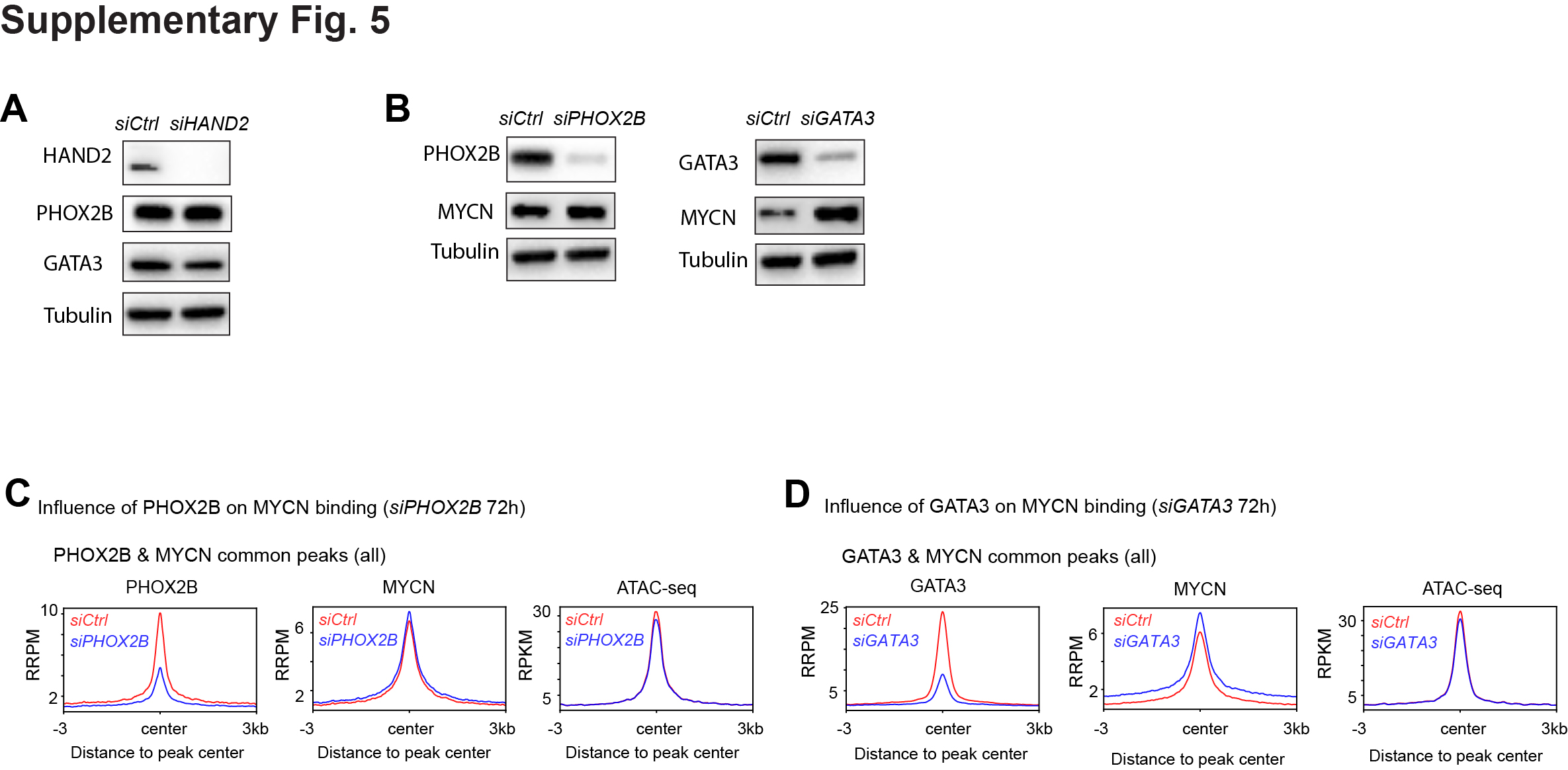

**Figure S5 (Extension of Figure 5). PHOX2B and GATA3 are not required for MYCN to bind to DNA.** (A) The effect of the knockdown of *HAND2* on the expression of PHOX2B and GATA3 at the protein levels shown by western blot. (B) The effect of the knockdown of *PHOX2B* or *GATA3* on the expression of MYCN at the protein levels shown by western blot. (C) Composite profile of ChIP-seq data shows the influence of the knockdown of *PHOX2B* on the average ChIP-seq signal of MYCN and the average ATAC-seq signal at PHOX2B and MYCN overlapped binding sites. Here all the PHOX2B & MYCN overlapped peaks are included. (D) Composite profile of ChIP-seq data shows the influence of the knockdown of *GATA3* on the average ChIP-seq signal of MYCN and the average ATAC-seq signal at these sites. Here all the GATA3 & MYCN overlapped peaks are included. RRPM: spike-in normalized, reference-adjusted reads per million mapped reads; RPKM: reads per kilobase per million mapped reads.

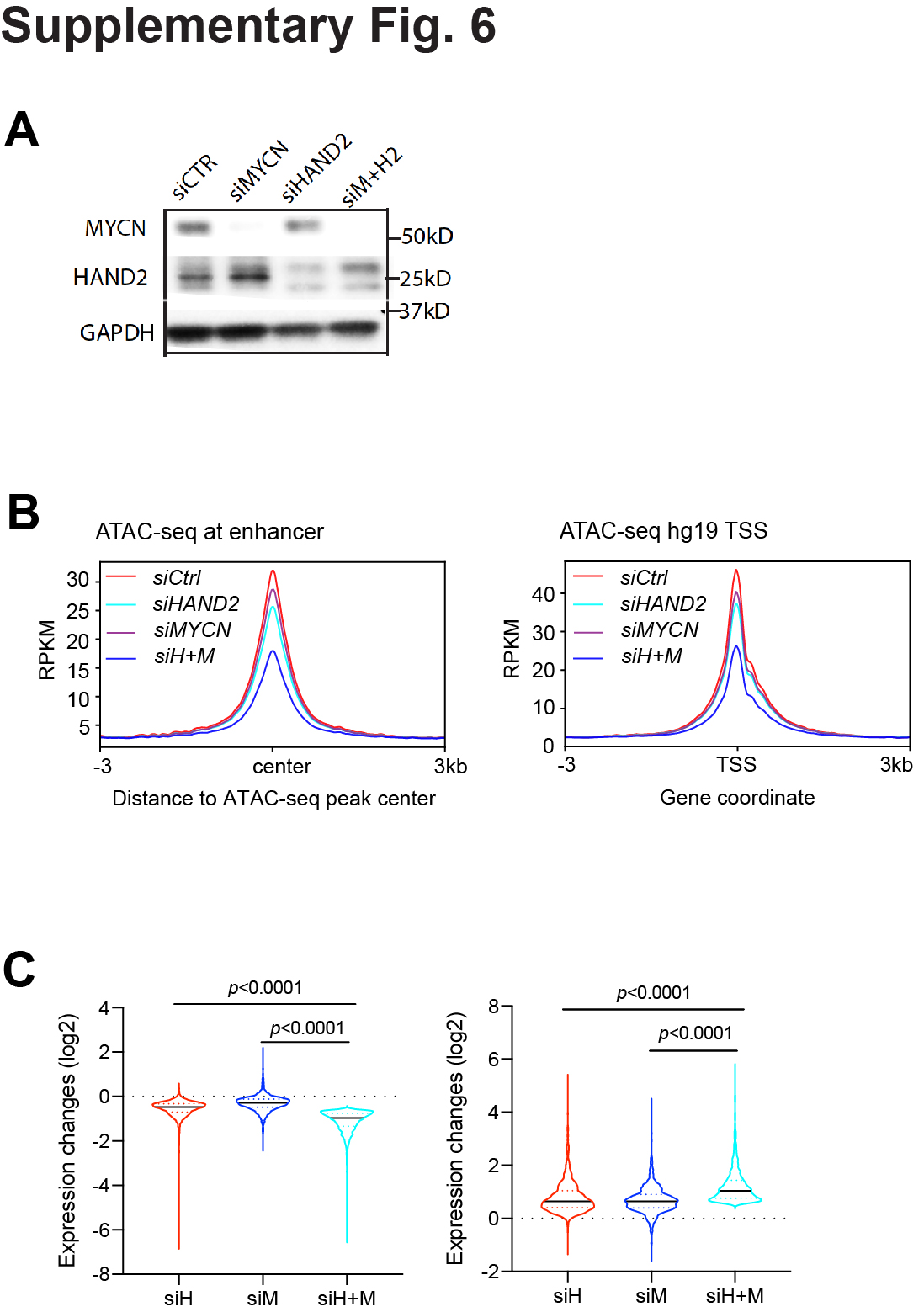

**Figure S6 (Extension of Figure 6). HAND2 and MYCN cooperatively regulate chromatin accessibility and gene transcription.** (A) Western blot assay shows that the silencing of *MYCN, HAND2* or both results in a decrease of their expression at protein levels. (B) Composite profile of ATAC-seq data shows a decreased average ATAC-seq signal at the active enhancer region when both *HAND2* and *MYCN* were knocked down (*siH+M*, left panel) (ATAC-seq peaks that overlapped with H3K27ac peaks and 1 kb away from the TSS). A decrease of average ATAC-seq signal is also observed when focused on all TSS sites when both *HAND2* and *MYCN* are knocked down in IMR32 cells for 72 hr (*siH+M*, right panel). (C) Relative gene expression changes at the mRNA levels after the knockdown of *HAND2, MYCN* or both shows a greater down-regulation (left panel) or up-regulation (right panel) of the gene expression when both *HAND2* and *MYCN* are knocked down than knocking down each one alone. The same gene lists used to generate the heatmaps from Fig.6b are used here. RPKM: reads per kilobase per million mapped reads. The *p*-value indicated is calculated in one-way ANOVA.

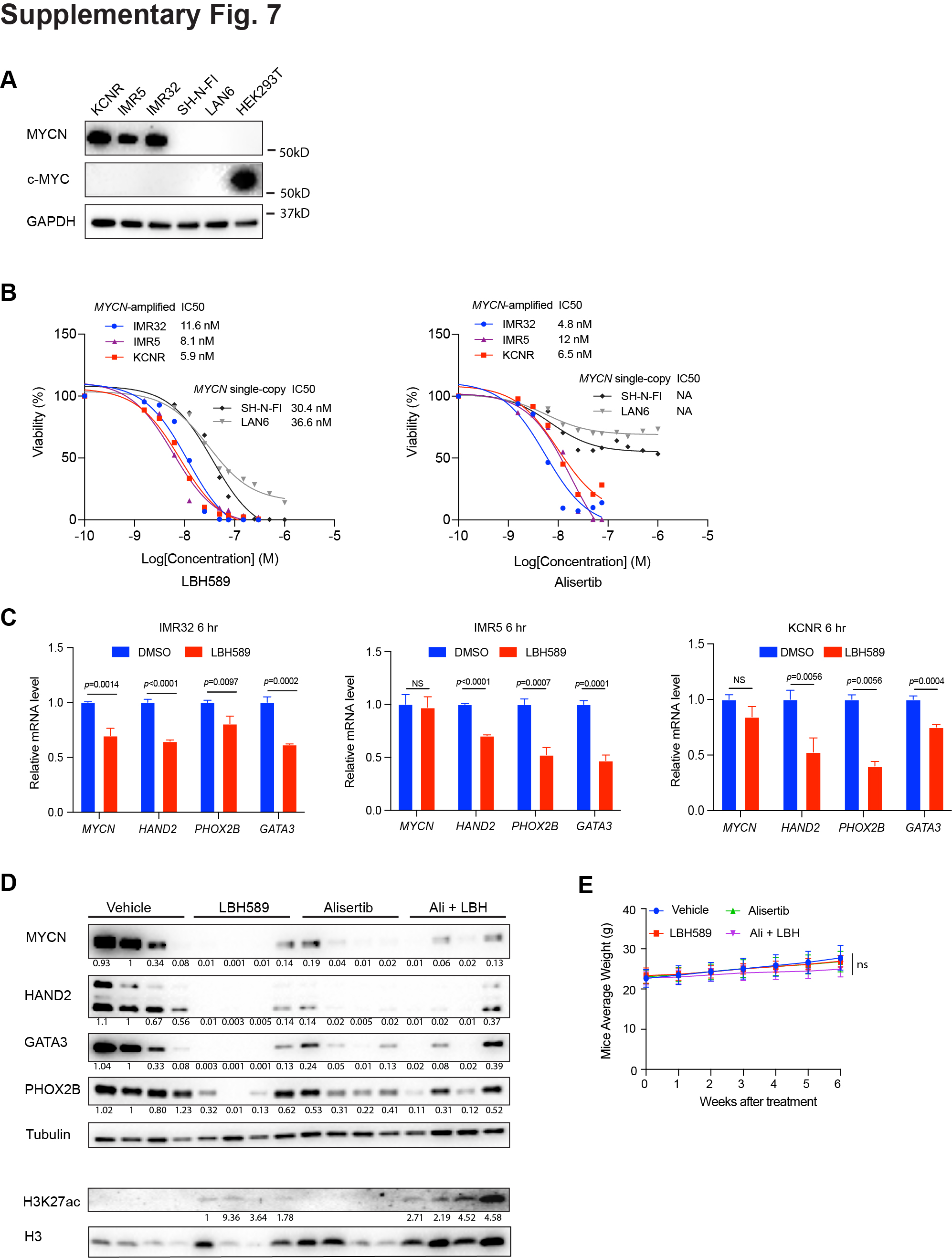

**Figure S7 (Extension of Figure 7). Targeting both MYCN and CRC TFs.** (A) The expression of MYCN and c-MYC at protein levels in NB cell lines and HEK293T cells are shown by western blot. HEK293T sample is a control to show c-MYC expression. (B) Dose-response curves of MYCN-amplified and MYCN single-copy NB cell lines after the treatment of LBH589 or alisertib for 72 hr. Percent cell viability relative to that of DMSO-treated cells is calculated based on CellTiter-Glo Cell Viability Assay. (C) The effect of the HADC inhibitor LBH589 treatment (6 hr) on *HAND2*, *PHOX2B*, *GATA3* and *MYCN* expression at mRNA levels in NB cell lines is shown by realtime PCR assay. The *p*-value indicated is calculated in Student’s t-Test. (D) The change of MYCN, HAND2, GATA3, PHOX2B at protein levels is observed in tumor samples collected from different group of mice that treated with vehicle or indicated drugs for 28 hr. The H3K27ac levels was first normalized to histone H3, then compared with H3K27ac level in lane 5; while the rest proteins were first normalized to tubulin, then compared with protein levels of each gene in lane 2. (E) The effect of the drug treatment on the body weight change of the mice.
